# Aligning transformer circuit mechanisms to neural representations in relational reasoning

**DOI:** 10.1101/2025.10.29.685457

**Authors:** Luke J. Hearne, Conor Robinson, Luca Cocchi, Takuya Ito

## Abstract

Relational reasoning—the capacity to understand how elements relate to one another—is a defining feature of human intelligence, yet its computational basis remains unclear. Here, we combined human neuroimaging (7T fMRI) with artificial neural network modeling to identify circuit-level analogues of human reasoning computations. Using the Latin Square Task, we found that humans and transformers were able to generalize the task reliably, while standard architectures used in cognitive neuroscience could not. Analyzing the transformer components revealed distinct computational roles: positional encoding captured the spatial structure of the task and aligned with representations in visual cortex, whereas attention encoded relational structure and mapped onto frontoparietal and default-mode networks. Attention weights tracked the relational complexity of the task, providing a computational analogue of reasoning demands. These results advance knowledge on the core algorithmic computations supporting complex reasoning, highlighting attention-based architectures as powerful models for investigating the neural and computational basis of higher cognition.

## Introduction

Relational reasoning is the ability to understand how elements interact to come to a conclusion, decision, or achieve a goal^1,2^. This capacity underpins a plethora of complex thinking skills that contribute to general intelligence^3–5^, including understanding analogy, inferring hidden causes, and generalizing patterns to novel contexts. In contrast to other animal species, humans are uniquely gifted at relational reasoning due to our increased capacity in processing multiple relations in parallel^1,6^. Although neuroimaging has mapped the neural correlates of relational reasoning implicating both frontal and parietal cortices^7–11^, progress toward understanding the underlying computations has been limited, partly due to the challenge of modeling complex intellectual processes^12^.

Modern artificial neural networks have demonstrated impressive performance on a range of complex tasks^13^, including forms of reasoning^14–16^. When paired with well-designed cognitive tasks and robust evaluations, these models can help reveal the computational mechanisms underlying specific cognitive processes^17^. When further combined with neural data, these models can adjudicate the similarities and differences between biological and computational implementations of these processes^18,19^. Using this approach, several studies have elucidated the computational mechanisms related to sensory processing^20^, memory and navigation^21,22^, and context-dependent decision making^17,23–25^, among others. However, advances in understanding the circuit mechanisms of higher-order cognitive processes, such as relational reasoning, remain limited. This is largely because designing reliable experiments to probe relational reasoning in both humans and ANNs is challenging, and commonly used neural architectures in neuroscience, such as recurrent (RNNs) and convolutional neural networks (CNNs), often fail to perform well on such complex reasoning tasks, limiting their utility as models of complex cognitive function.

Recent advances in artificial intelligence have been driven by the computational power of the transformer architecture^13^. In brief, transformers combine three key neural network components: positional encoding (PE), the attention mechanism, and a standard multilayer perceptron (MLP). Similar to how retinal cells receive 2D information from the visual field while maintaining the correct spatial organization of a stimulus, PE preserves the order and arrangement of input tokens (e.g., a spatial receptive field), providing a useful inductive bias for organizing inputs^26^. On the other hand, *attention* allows each element of an input (e.g., each word in a sentence) to query all other elements and decide which ones are relevant with respect to their positioning^27^. This allows the model to understand relationships between elements in the context of the entire input. This specific mechanism provides a computational link to the flexible coordination required to relate elements to one another in relational reasoning. Finally, the MLP portion of the transformer – a standard neural architecture – appears to behave as a key-value memory store^28^. While not strictly brain inspired, computational parallels have been drawn between transformers and cortico-thalamic circuits^29^, neural visual information routing^30^, and models of the hippocampus^31^. For example, transformers can reproduce empirical spatial representations from hippocampal place and grid cells^31^. Nevertheless, exactly how transformers, or any computational model, might implement relational reasoning computations remains unclear, especially given their limited application in cognitive science using well-controlled psychological experiments.

Motivated by this gap, we investigated the computational mechanisms of relational reasoning across various artificial neural networks and compared them to human performance and brain imaging data on an established relational reasoning task: the Latin Square Task (LST)^32^. The LST is a nonverbal spatial relational reasoning task that leverages relational complexity theory^1^ to systematically manipulate reasoning difficulty. We found that, while humans reliably solved LST puzzles, both standard MLP and recurrent neural network architectures failed to generalize beyond the training set. In contrast, transformer models were capable of robust generalization. Despite not being biologically inspired, the transformer’s PE and attention mechanisms captured computational strategies that paralleled human neurocognitive processes. Specifically, PE captured the spatial structure of the task and aligned with representations in visual cortex, whereas attention tracked the relational complexity of the task and mapped onto frontoparietal and default-mode networks. Moreover, fine-grained analysis of the attention weights suggested that greater attentional ‘spread’ (i.e., the density of the weights) was associated with more complex reasoning problems, providing a computational analogue of the relational integration demands of the task. Finally, we illustrate how different layers of transformer representations produce heterogeneous representations that organize from lower-level to higher-order cortical representations. Together, these findings demonstrate that transformer neural networks can capture key computational motifs of cortical systems during complex reasoning.

## Results

### The Latin Square Task in humans and neural networks

We start by contrasting performance between humans and neural networks on the LST^32^. The LST was developed using relational complexity theory^1^ to probe relational reasoning in humans. Within this framework, processing demand scales with the number of relations that must be considered simultaneously in a given problem. This contrasts with, for example, working memory tasks, wherein the number of items to be kept in mind dictates the difficulty. Working memory is considered the ‘workspace’ in which relational representations are constructed; it is necessary but not sufficient for the integration step that defines relational reasoning^33,34^.The behavioral performance associated with increments in relational complexity on the LST, and their relationship to independent measures of fluid intelligence are well established^35,36^.

The task involves inferring missing elements in a grid based on relational constraints, analogous to the game Sudoku, which has become a popular and challenging benchmark for AI systems^37^. Specifically, participants are shown a 4×4 grid populated with shapes, blank spaces, and a target question mark. Participants are asked to determine the shape that fits in the target cell, ensuring that each shape only appears once in each row and column. Puzzles can be organized into three increasingly complex reasoning conditions^1^ that are dictated by the number of vectors that need to be taken into consideration to solve each given puzzle (an example of each is shown in **Fig. 1A**). Importantly, the rule structure, stimuli, grid format, and response demands are held constant across conditions, isolating increases in relational integration from other working-memory demands. Thus, the LST offers a tractable, yet meaningful benchmark for probing the computational mechanisms underlying relational reasoning in both humans and machines, while also being well-suited for measuring human brain activity during task performance.

**Fig. 1.**
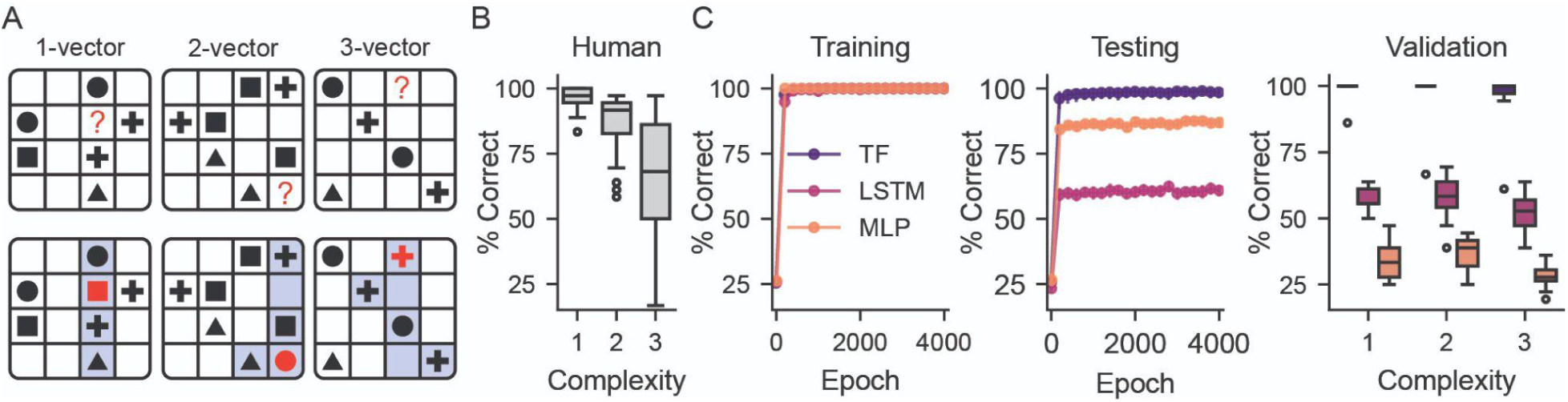
Experiment setup and model performance. **A.** Example puzzles from the Latin Square Task (LST) for 1-, 2- and 3-vector reasoning complexity conditions. Blue highlights in the bottom panel indicate the cells and vectors which contain the required information to solve each problem **B.** Human (N_participants_=40) performance on the task (left). **C.** Average ANN (N_seeds_=15) task accuracy in the training (left), testing (middle) datasets and validation (right; identical problems to brain imaging data). Boxplots show the median and interquartile range; whiskers extend to 1.5× the interquartile range and points beyond plotted as outliers. TF; Transformer, LSTM; Long Short-Term Memory, MLP; Multilayer perceptron.

As expected, human performance on the LST decreased as relational complexity increased (*F*(2,78)=55.29, p<.001, *η^2^_G_* =.43, **Fig. 1B**). Follow up *t*-tests confirmed the direction of this effect, 1-vector puzzles (*M*=96.67%, *SD*=4.13%) were solved more often than 2-vector puzzles (*M*=87.01%, *SD*=11.01%, *t*(39)=6.78, *p*< .001, *d=1.15)* and 2-vector puzzles were solved more often than 3-vector puzzles (*M*=66.25%, *SD*=22.46%, *t*(39)=5.94, *p*< .001, *d=1.16)*. Accuracy in the most difficult condition (3-vector) correlated with performance on a fluid intelligence test (The Raven’s Advanced Progressive Matrices, Spearman’s ρ =.48, p=.002).

We compared three neural network models: an encoder-only Transformer, a Long Short-Term Memory (LSTM) network, and a multilayer perceptron (MLP, see **Methods** for details). These comparison models were included to demonstrate that with no additional architectural changes, the transformer, as opposed to vanilla MLP and LSTM ANNs, is well-suited to solving the LST. All three models achieved perfect accuracy in the task in the training dataset (8000 puzzles, **Fig. 1C**). However, in 1000 unseen test puzzles drawn from the same distribution as the training data, model performance dropped substantially for the non-Transformer ANNs (mean performance; Transformer=98.51%, LSTM=60.89%, MLP=86.91%). Finally, the ANNs were tested on the same puzzles that humans completed. These puzzles were designed with low correlation with the training dataset (jaccard dissimilarity > 0.8), thus representing a difficult out-of-distribution generalization task. While all models performed above chance (chance=25%, mean performance; Transformer=97.65%, LSTM=55.49%, MLP=33.02%), only the Transformer could successfully generalize and solve the task at a level comparable, or exceeding, human performance. Main effects of relational complexity were also observed in the transformer (see **Supplementary Material**).

### Mapping architectural components of transformers to neurocognitive mechanisms

Relational reasoning in humans depends upon cortical systems that encode structures and integrate relational information. Converging evidence from lesion, neuroimaging and brain stimulation studies have consistently implicated a frontoparietal network (FPN) of brain regions as central to relational reasoning^2,7,8,38–41^. To investigate whether analogous computations emerge in the neural network model, we examined how distinct components of the transformer architecture correspond to these functions. In particular, we evaluated the two architectural components that are unique to the transformer model; the positional encoding (PE) and self-attention mechanism (**Fig. 2A**). PE is a transformer mechanism that provides spatial information about the location of tokens. Since transformers process tokens in parallel rather than sequentially, in the absence of PE, the transformer would not know the correct location of each symbol in the LST. Since the LST is organized as a 2D grid, we utilized a PE that preserved the pairwise distance between tokens in 2D. Strictly speaking, while the 2D arrangement of the LST is not required to solve the problem – most state-of-the-art solvers on LST-style tasks actually use a 1D string of codes as input^42^ – we hypothesized that a 2D structure would provide a helpful spatial scaffold from which to perform the LST (**Fig. 2B**).

**Fig. 2.**
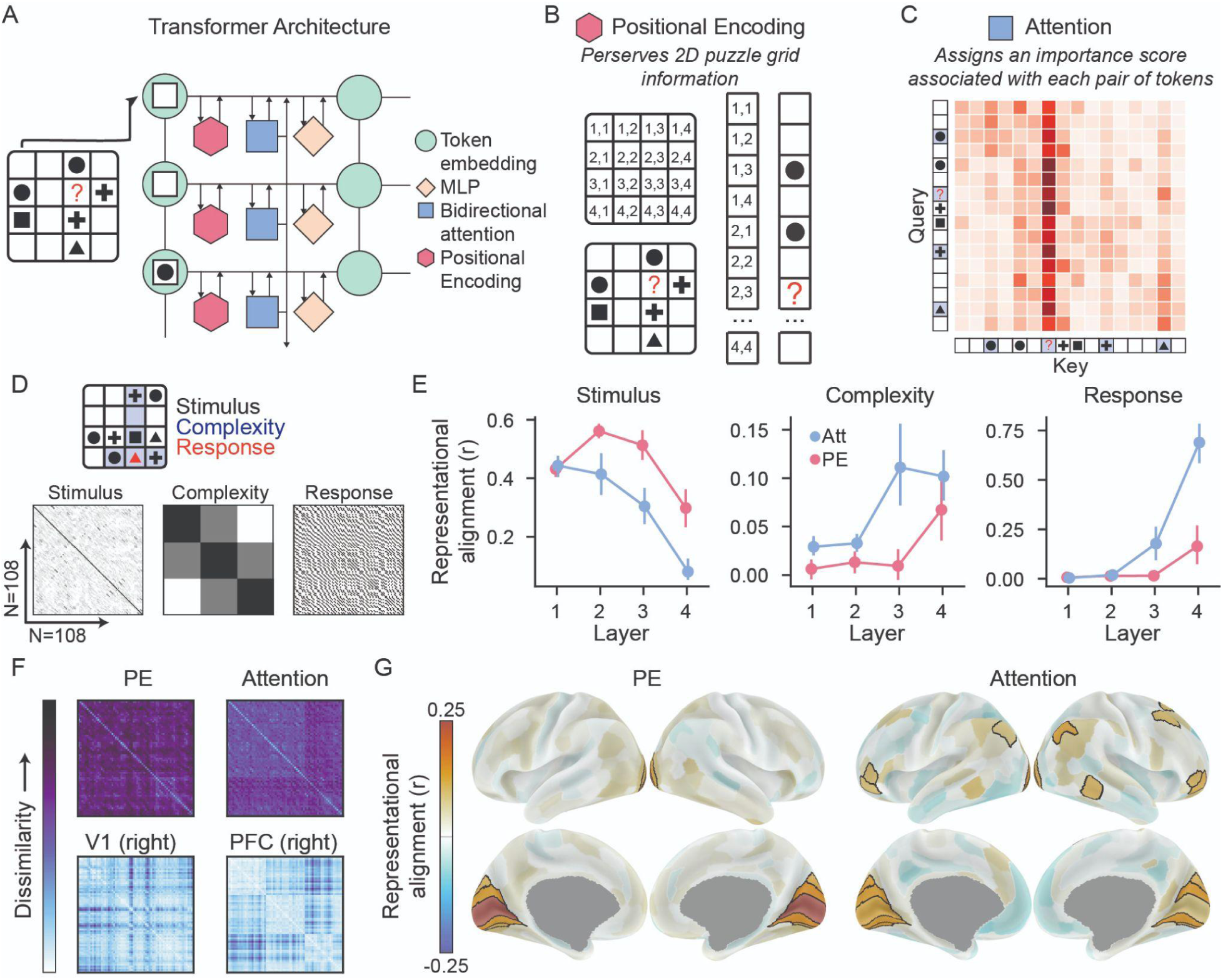
Transformer circuit components and their alignment with task structure and neural representations. **A.** Schematic of the transformer architecture for a single layer. **B.** Schematic of the positional encoding (PE) mechanism. PE information describes the two-dimensional organisation of the puzzle, and therefore the critical rule that each shape can only appear once in every row or column. **C.** Attention weights (*QK^T^* matrix) for the example 1-vector puzzle. Attention weights quantify how strongly each token (puzzle element) relates to all other tokens, guiding the model’s focus during information processing (darker red colors indicate stronger weights). For example, the column of stronger weights associated with the target (“?”) indicates a disproportionate amount of information being broadcast to other puzzle elements. **D.** Experimenter defined models of stimulus, task complexity and response. **E.** Representational alignment between task feature models and transformer post-PE and - Attention mechanisms. Error bars denote 95% bootstrapped confidence intervals. **F.** Example representational dissimilarity matrices derived from the transformer (top row) and fMRI data (bottom row). **G.** Representational alignment between brain and transformer (PE and Attention) embeddings. Black borders indicate significant ANN-fMRI representational alignment (permutation and FDR corrected).

Once the geometry of the tokens (i.e., 2D grid) is accurately processed, the token representation (along with the tag of its location) is sent to the downstream self-attention. There, the self-attention mechanism functions like a dynamic spotlight of visual attention; it evaluates every token’s relevance to every other token, assigning “importance” weights to each token-pair. For example, in natural language processing PE aims to embed the order of each word in a sentence, whereas attention aims to identify the relevant relationship *between* words to derive meaning. In the LST, attention is the natural candidate to capture the relational structure needed to solve the task. More concretely, this “spotlight of visual attention” is governed by the dot product of two vectors, for every pair of tokens: the queries (*Q*; the receiver) and keys (*K*; the sender) vector (**Fig. 2C** shows an example *QK^T^* matrix, or attention weights). Functionally, the *Q* vector sends a “query” asking for a specific type of information, and the *K* vector provides information to the query. The dot product then provides an indication of how well the *Q* and *K* vectors matched. In the example provided in **Fig. 2C**, we can see that all tokens received information regarding the target “?”, though information routing was disproportionately sent to tokens in the row and column of the probe. Attention weights are then applied to the value (*V*) vectors, which encode the information content of each token, producing an output that reflects a weighted integration of routed signals (*QKV*) that is added to the transformer’s residual stream.

Next, we examined the influence of the transformer’s architectural mechanisms on model representations. To quantify representational content at different stages of processing, we used representational similarity analysis (RSA)^43,44^, which enables direct comparison between the similarity structure of model embeddings and experimenter-defined models of task-relevant features. For each transformer layer (1-4), and component (PE, attention) we computed representational dissimilarity matrices (RDMs) across the 108 test puzzles using layer-wise token embeddings. For readability, we refer to the positionally-informed input representation as PE (*e*(*x_i_*) + *p_i_*), and the post-attention representation as ‘attention’ (*X_PE_* + *softmax*(*QK^T^*)*V)*.

Three experimenter models were contrasted; *stimulus*, the similarity between input puzzle features; *complexity*, the relational complexity of the puzzles (defined by the number of vectors needed to solve the puzzle); and *response*, the similarity between correct puzzle solutions (**Fig. 2D**). We refer to the resulting similarity as representational alignment^45^. Importantly, as opposed to stimulus and response information which are the input and output of the model, relational complexity is an emergent property of the task structure. Critically, the relational complexity model did not correlate with either the stimulus or response models (see **Methods**). Given the distinct computational roles of these components, we expected PE representations to align most strongly with stimulus features, whereas attention representations would align more closely with relational complexity and response-related experimenter models.

We observed a main effect of architectural feature in all three experimenter models; PE demonstrated higher similarity to input stimulus (M_PE_=0.45, M_ATTN_=0.31, *F*(1, 14)=575.27, *p*<.001, *η^2^_G_* =.37), whereas attention activations showed higher similarity to reasoning complexity (M_PE_ =0.02, M_ATTN_ =0.07, *F*(1, 14)=131.59, *p*<.001, *η^2^_G_* =.20) and response models (M_PE_ =0.05, M_ATTN_ =0.22, *F*(1, 14)=144.92, *p*<.001, *η^2^_G_* =.38) (**Fig. 2E**). Main effects of layers, and interactions between layer and architectural features were also observed (see **Supplementary Table 1**). This analysis confirms the role of PE in organizing the visual geometry of the puzzle, and the attention mechanism in relational processing and response generation. Consistent with recent mechanistic accounts of transformers^28,46^, response-related information observable in the attention circuit reflects how the model integrates and applies routed information in the grid, whereas the MLP encodes the semantics of the routed information.

We next sought to assess whether similar patterns of representation emerged in human brain activity and each transformer architectural component when averaging across layers (**Fig. 2F**). For the neural data, RDMs were computed across the 108 test puzzles for each brain region (N=360), using vertices belonging to each region. We observed significant alignment of representations between specific regions of the primary visual cortex and PE representations (**Fig. 2G**, *p*_FDR_ < .05**, Supplementary Table 4**). For embeddings processed after the attention mechanism, significant alignment was additionally observed within multimodal brain regions belonging to frontoparietal and default-mode networks (*p*_FDR_ < .05, network affiliations were defined as in^47^). Compared with the upper noise ceiling bound (95th percentile across 1000 split-halves), on average the models accounted for 26.22% (PE) and 23.08% (Attention) of the explainable variance in the fMRI data (see **Supplementary Fig. 3**). These regions substantially overlap with areas previously implicated in distinct components of relational reasoning, including anterior prefrontal cortex in integrating multiple relations and posterior parietal cortex in representing relational structure^39,48,49^. These results provide evidence that cognitively meaningful representations – PEs with stimulus-related features, and attention with relational complexity-related features – converge across transformers and human brains, despite substantial architectural differences. We verified in control analyses that the explained variance between the transformer model embeddings, experimenter-defined models, and the fMRI data is largely attributed to the model’s learned representations (**Supplementary Figs. 1-2, Supplementary Tables 2-3**).

### Importance of positional encoding on learning meaningful stimulus features

The transformer’s PE mechanism produced representations that aligned with the primary visual cortex and stimulus information (**Fig. 2E-G**). Architecturally, the 2D PE introduces a useful inductive bias, enabling the transformer to capture token order (i.e., the puzzle shape), an essential feature for LST performance. To test whether this bias was necessary, we removed the 2D structure and replaced it with a learnable vector embedding initialized at random such that every pair of tokens (or grid element) was equidistant at initialization. This provided a fairer comparison with the MLP and LSTM models, which similarly do not encode positional biases across tokens. Recent work has demonstrated that varying parameter initializations can lead to significant differences in the structure of internal representations^50,51^. In particular, weight initializations sampled from a large distribution (e.g., *N*(0, σ) for large σ) tended to produce random, high-dimensional representations (lazy learning), while weight initializations sampled from a small distribution (small σ) tended to learn structured, interpretable representations (rich learning). Within the context of PE for transformers, we initialized learnable vector embeddings with rich (small σ) and lazy (large σ) initializations to assess generalization performance on the out-of-distribution validation set. We also subsequently inspected the alignment between the learned PE vector representations and the brain (**Fig. 3A**)^24^. This variation in PE is specific to the current results; fixed 2D PE was used for all other analyses.

**Fig. 3.**
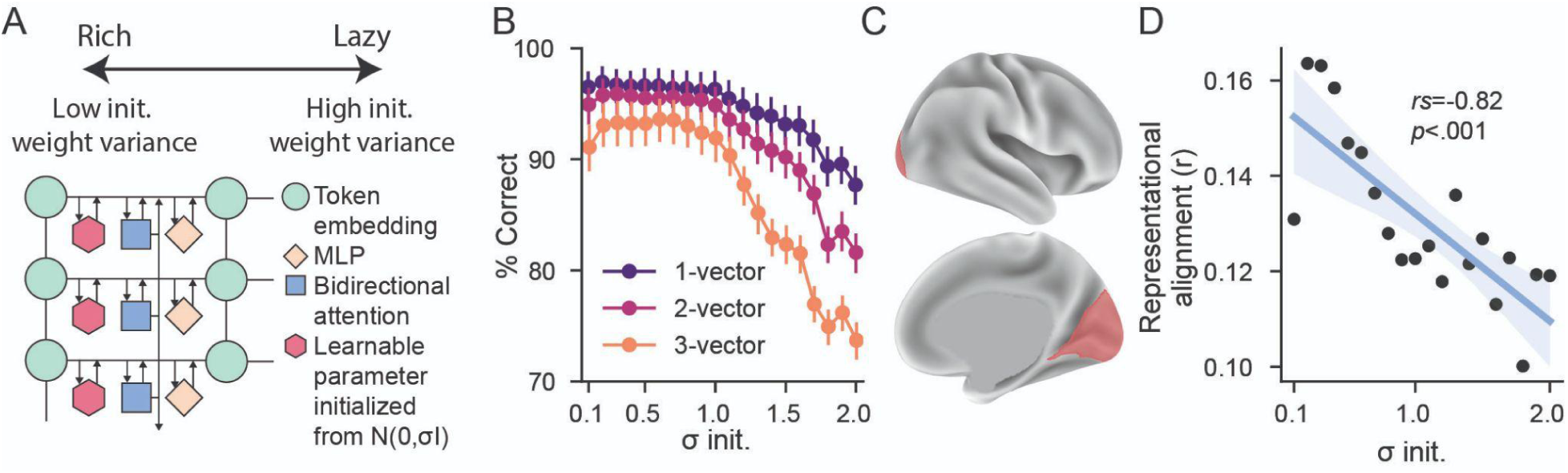
Influence of rich and lazy positional encoding weight initialization on generalization performance and brain-transformer representational alignment. **A.** Schematic of transformer architecture with learnable PE parameters initialized from rich and lazy regimes. **B**. Task generalization performance when positional encoding (PE) weight initializations (init.) are systematically altered from rich (small σ) to lazy (large σ). Error bars denote 95% bootstrapped confidence intervals. **C.** Regions of interest derived from the prior fMRI PE RSA analysis. **D.** Representational alignment between brain and PE within regions of interest across PE initializations.

We found that transformer task accuracy in the out-of-distribution validation set was related to initialization parameters, such that low σ (i.e., rich) models performed significantly better than high σ (lazy) models (F(19, 266)=65.93, p<.001, *η^2^_G_* =.73, **Fig. 3B**). Similar to the human data, there was also a main effect of reasoning complexity (F(2, 28)=65.71, p<.001, *η^2^_G_* =.07), and a weak interaction effect between weight initialization and reasoning complexity (F(38, 532)=5.85, p<.001, *η^2^_G_* =.06). The complexity-initialization interaction suggests that “lazy” models had particularly poor performance in the most complex reasoning condition.

Next, to better understand why richly learned PEs exhibited better generalization performance, we sought to characterize the learned PE representation. In the previous section, we demonstrated that PE representations that were derived from 2D encodings (i.e., those that preserved the original 2D grid without learning) exhibited high representational alignment with human visual cortex representations (**Fig. 2G**). Thus, we hypothesized that models with *richly learned* PEs (i.e., the models that exhibited improved generalization performance) would learn structured PE representations that had greater alignment with human visual cortex representations. We therefore evaluated whether alignment across fMRI data and transformers was related to the initialization parameter.

Using brain regions of interest defined in the prior analysis (outlined regions in **Fig. 2G** related to PE; **Fig. 3C**) we observed that low σ PE initialization demonstrated higher brain-transformer alignment, specifically in visual regions (*rs*=-.82, p<.001, **Fig. 3D**). These results underscore the value of learning structured visual representations (through the PE) for robust relational reasoning, and show again that transformers – despite their architectural differences – can learn meaningful cognitive and brain-like representations.

### The computational relevance of attention weights in relational complexity

Our earlier analysis illustrated that the transformer’s attention mechanism produced embeddings sensitive to relational complexity using RSA (**Fig. 2E**). This fits with the architecture, as the attention circuit evaluates the importance of every pairwise interaction between tokens (i.e., elements in the LST grid). Similarly, in the LST, greater relational complexity demand is operationalized as an increase in the number of vectors (1-, 2-, or 3-vectors) needed to be integrated simultaneously to solve a puzzle^1^. Thus, we expected lower-complexity puzzles to exhibit a more localized relevance structure, whereas higher-complexity puzzles would require coordination across a broader set of interdependent elements. Accordingly, the *QK^T^* matrix, or *attention weights*, which describe the mappings between tokens (16 x 16 for each puzzle, examples shown in **Fig. 4A**), provide a window into how relational complexity might be organized in the transformer. Specifically, each element in the *QK^T^* matrix reflects how similar the query and key vectors are for that token pair. The transformer uses these values differently: A strong *query* value for a token reflects how a token receives information, whereas a high *key* value for a token reflects how widely broadcast that information is to all other tokens.

**Fig. 4.**
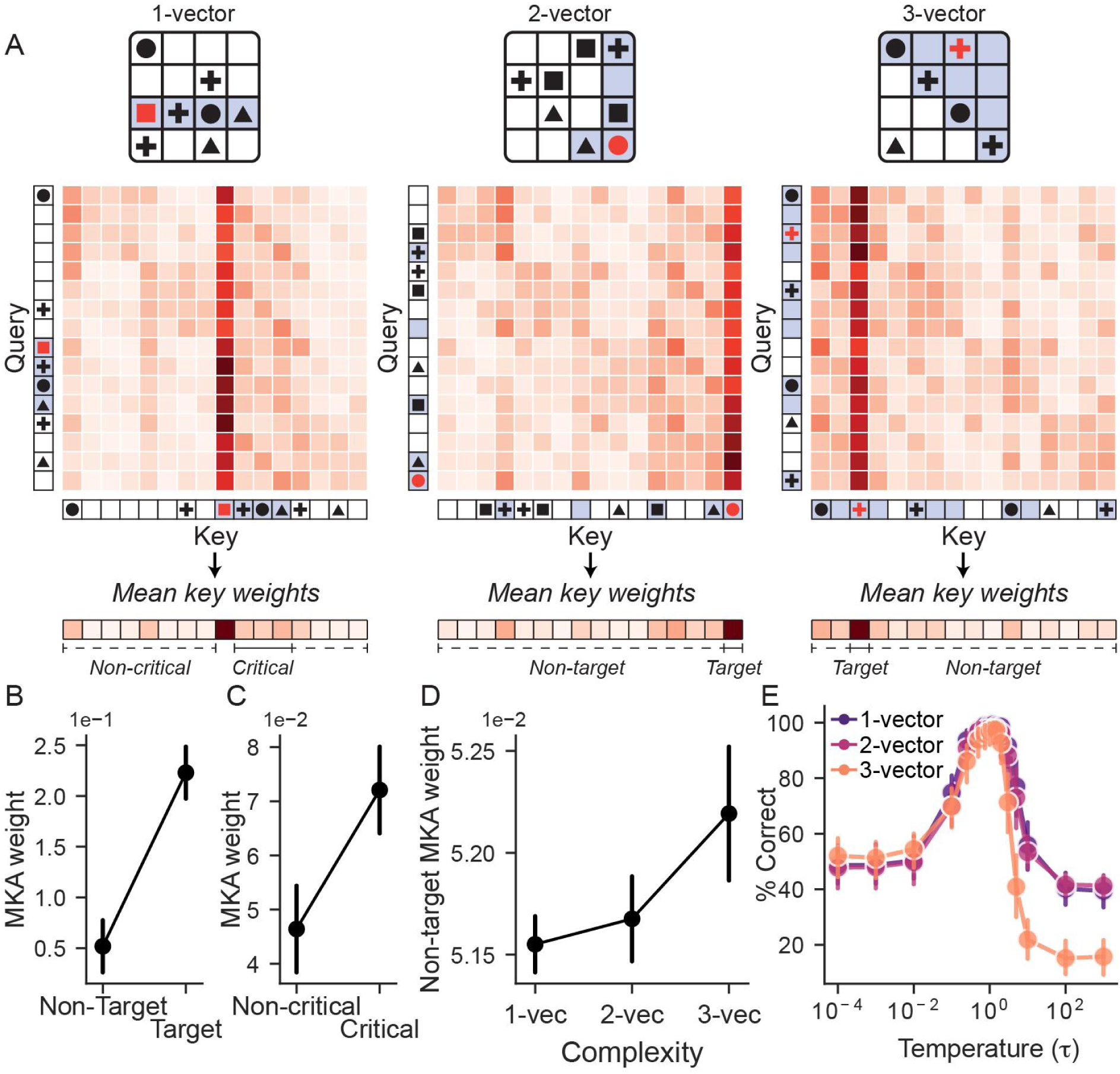
Relational complexity in the transformer’s attention mechanism. **A.** Example puzzles from each relational complexity condition and associated attention weight (*QK^T^*) matrices. The attention weight matrices denote the relationship between each of the 16 puzzle elements, averaged across layers and transformer model initializations. We calculated mean *key* attention weights, which represent the extent to which each puzzle element (or token) “broadcasts” its information to other puzzle elements. **B.** Mean key attention (MKA) weights for the target compared to non-target puzzle elements. **C.** Comparison of MKA weights between critical and non-critical cells in one-vector problems. In the example shown, only the three cells of the third row are critical for solving the problem. **D.** Non-target MKA weights as a function of relational complexity. Error bars for panels **B-D** represent within-subject 95% confidence intervals (Cousineau–Morey corrected for repeated-measures designs). **E.** Task generalization performance when the temperature parameter (controlling the entropy of the attention weight distribution per row) was systematically manipulated from low (10^-4^) to high (10^3^). Points represent group means for each relational complexity condition and error bars denote 95% bootstrapped confidence intervals.

There are several testable hypotheses that lend themselves to interpreting the self-attention mechanism in the LST. We start by focusing on the attention weight of the target (“?”) element (equivalent to the red symbols in **Fig. 4A**). We quantified this by deriving each puzzle element’s mean *key* weight (averaging across queries; column sum). This value reflects how widely broadcast each token’s information is to all other puzzle elements. We found that the mean key weight was higher for the target when contrasted with other non-target puzzle elements (*t*(14)=8.95, *p*<.001, *d*=3.48, **Fig. 4B**). This result is intuitive, as knowledge of the target location is crucial to solving the puzzle.

Next, we tested whether the correct strategy needed to solve one-vector problems could be related to attention weights. Specifically, one-vector problems can *always* be correctly solved by integrating information in the appropriate column *or* row. This means that the transformer’s attention mechanism only needs to attend to elements in either a row or column. For example, the 1-vector problem in **Fig. 4A** requires only integration of elements in the third row (cross, circle, triangle, therefore target=square). We found that ‘critical’ cells have higher mean key attention weights than non-critical cells (*t*(14)=4.30, *p*<.001, *d*=1.69, **Fig. 4C**), suggesting the model appropriately learns to broadcast information from critical cells to the target.

Finally, we examined how attention weights varied with increasing reasoning complexity (**Fig. 4D**). As higher-complexity puzzles require information to be integrated across a larger number of grid elements, we hypothesised that complexity would be associated with more distributed information broadcasting from non-target elements (i.e., grid elements that determine the correct symbol in the target’s location). To test this, we quantified non-target average key weights as a function of relational complexity. Unlike the earlier analyses (where we focused on how the *target* broadcasts information), this analysis assessed how broadly *non*-*target* elements broadcast information as complexity increases, providing a complementary view of how attention weights are shaped by relational demands. Consistent with this intuition, we found that attention weights of non-target elements increased as reasoning complexity increased (χ^2^(2)=13.73, Kendall’s *W*=0.46, *p*<.001), which was largely driven by increased non-target key weights in 3-vector problems (1-vector vs. 2-vector; *t*(14)=-2.22, *d*=0.03, *p_Holm_*=.082, 1-vector vs. 3-vector; *t*(14)=-3.25, *d*=0.14, *p_Holm_*=.017, 2-vector vs. 3-vector; *t*(14)=-2.25, *d*=0.11, *p_Holm_*=.082). This analysis shows how transformers use attention weights to solve relational reasoning problems of varying complexity, offering a computational analogue of relational complexity within the transformer attention mechanism

We next modulated the attention weights by systematically varying the *temperature* (τ) parameter applied to the QK^T^ matrix in the attention mechanism (**Methods**). Temperature rescales the *QK^T^* logits without altering learned parameters, thereby controlling the entropy of the resulting attention weight distribution. Lower temperatures (τ < 1) produce sharper, lower-entropy distributions, whereas higher temperatures (τ > 1) yield progressively flatter, higher-entropy distributions. We evaluated a wide range of temperatures (τ = 0. 0001 to τ = 1000) to characterize the full regime of model behavior. Across temperatures, we assessed model accuracy on held-out puzzles and quantified model–human correspondence by correlating per-puzzle transformer accuracy (averaged across seeds) with human accuracy (averaged across participants).

Departures from the default temperature (τ=1), both low and high, were associated with reductions in model accuracy (**Fig. 4E**). On face value, this makes sense; very low temperatures would result in distributions unable to simultaneously attend to the multiple tokens needed to solve the task. Conversely, at very high temperatures, the attention weights become flatter and are seemingly unable to focus on the critical tokens needed to solve a given puzzle, as attention becomes dispersed. This analysis suggests a ‘goldilocks zone’ of appropriate attention weight distributions associated with good model performance. Notably, like humans, accuracy decrements at higher temperatures were most pronounced for 3-vector problems. Consistent with this observation, higher temperatures yielded the strongest correlation with human accuracy patterns across puzzles (peak correspondence at τ=3; Spearman’s ρ = 0.61, *p*_FDR_<.001, see **Supplementary Fig. 4** and **Supplementary Table 6**). Together, these findings indicate that modulating attention entropy systematically alters task performance, with higher-entropy regimes mirroring aspects of human relational integration errors.

### Brain-transformer representational alignment across layers

Hierarchical organization is a defining principle of cortical computation^52^. In cognition, low-level sensory representations are progressively transformed into increasingly abstract high-level representations in transmodal cortex. Prior work, particularly in visual processing and object recognition, has drawn parallels between hierarchical information processing observed in the brain and ANNs^20^. Our prior analysis (**Fig. 2E**) showed that transformer embeddings exhibited high similarity to stimulus features in early layers, which progressively decreased across layers. Conversely, similarity to relational complexity was initially low and increased across layers. Here we tested if layer-dependent representational alignment between the brain and transformer outputs mirrored cortical hierarchies. To do so, we computed the alignment between the fMRI RDMs (derived across the 108 test puzzles) to each layer of the transformers MLP embeddings.

The transformer-fMRI representational alignment was dynamic; appearing in visual cortex in early layers before progressing to parietal and frontal cortex in later layers (**Fig.5A**, *p*_FDR_ < 0.05, see **Supplementary Table 5**). When contrasted with the upper noise ceiling bound (95th percentile across 1000 permutations), on average, the model accounted for 19.43% (Layer 1), 35.14% (Layer 2), 42.10% (Layer 3), and 78.25% (Layer 4), of the explainable variance in the fMRI data. This stepwise increase in explained variance may reflect the experimental design, in which the fMRI task was constructed to elicit reasoning-related neural activity that is better captured by later, more abstract model layers. These estimates should nevertheless be interpreted cautiously, as they depend on the specific noise-ceiling estimation procedure used (see **Methods, Supplementary Fig.3**).

**Fig. 5.**
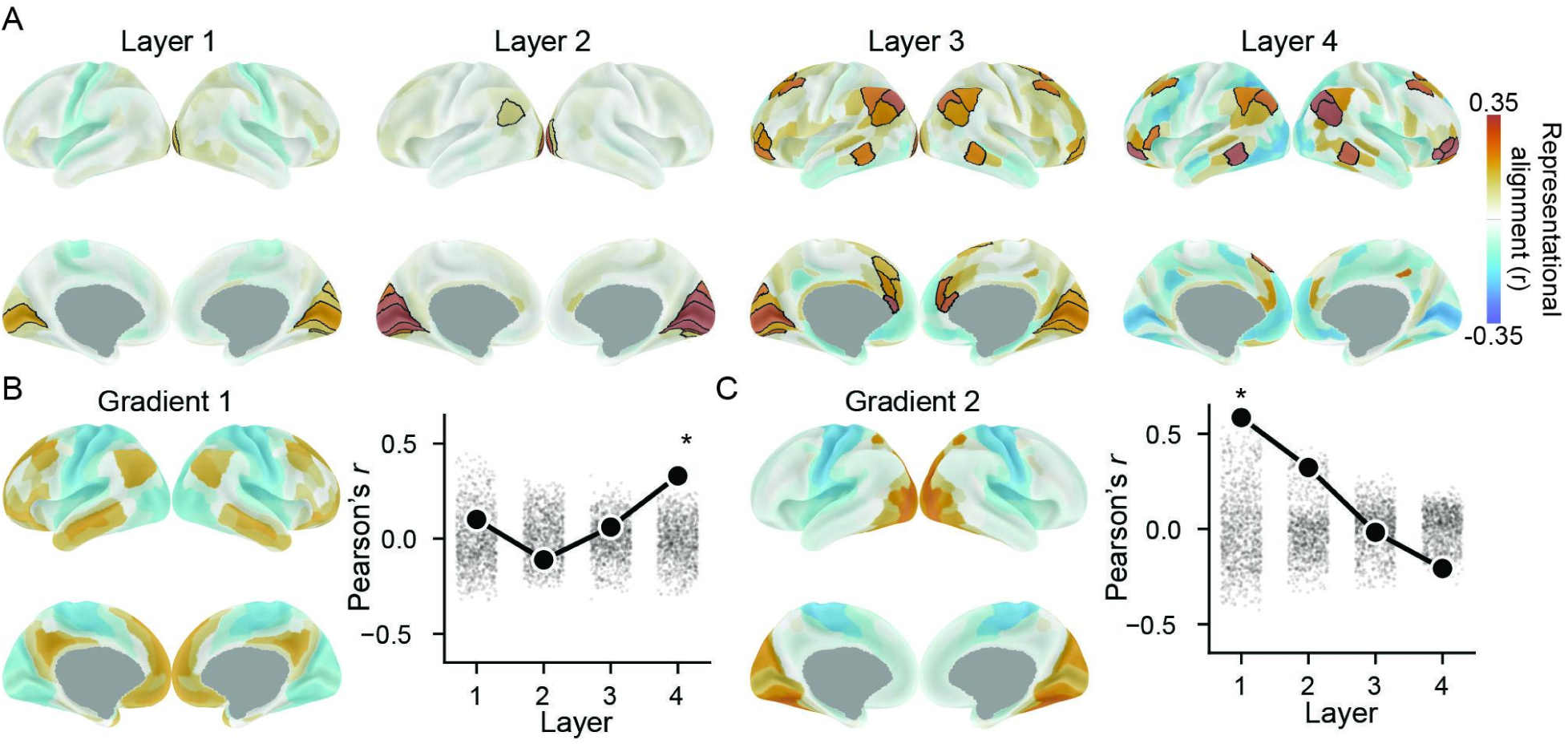
Cortical hierarchical alignment across Transformer layers. **A.** Layer-dependent representational alignment between transformer and human fMRI during relational reasoning. Each column denotes a layer in the transformer architecture. Black borders indicate significant ANN-fMRI representational alignment (permutation and false discovery rate corrected). **B.** Primary functional connectivity gradient defined by Margulies and colleagues^53^ and its correlation with transformer-fMRI representational alignment brain maps (shown in panel A). **C.** Secondary functional connectivity gradient and its correlation with transformer-fMRI representational alignment brain maps. Grey data show spin test permutation distributions. Asterisks denote spin test and FDR corrected significance (*p*<.05).

We next compared these alignment maps with the principal functional connectivity gradient, which captures a unimodal-to-transmodal cortical hierarchy, and the secondary connectivity gradient, which reflects a unimodal visuo-motor axis, as reported in^53,54^ (**Fig. 5B-C**). This analysis revealed significant correspondence between transformer–fMRI alignment patterns and the primary connectivity gradient in the final layer (*r=*0.33, *p_FDR_=*.009), as well as significant correspondence with the secondary gradient in the first layer (*r*=0.58, *p_FDR_*<.001). Together, these results indicate that representational transformations across transformer layers adhere to established cortical hierarchies.

## Discussion

We investigated the cognitive and computational mechanisms of relational reasoning. To do so, we analyzed representations of both ANNs and human task fMRI data during LST performance. While standard out-of-the-box ANN architectures used in computational neuroscience failed at generalizing the task (i.e., MLPs, LSTMs), transformers were able to generalize the LST reliably.

By isolating the representations produced by the transformer’s unique components – namely its PE and attention mechanism – we characterized the precise representations produced by the transformer for successful LST performance. Specifically, PE produced representations that were highly sensitive to the visual, grid structure of the LST puzzle. On the other hand, the attention mechanism produced representations that aligned with the relational complexity conditions of the LST. We further found that the representations produced by each architectural component could be similarly observed in the human brain; PE and attention representations were representationally aligned to neural systems responsible for visual and higher-order cognitive control representations, respectively. When focusing on inter-layer transformer representations rather than intra-layer architectural components, we found a systematic transition from lower-to higher-order neural representations across successive layers, mirroring hierarchical processing observed in other biological systems (e.g.,^20^). Our findings build an algorithmic account of both the computational mechanisms required to perform and generalize relational reasoning in the LST, and how they correspond to those in the human brain.

Prior studies in computational neuroscience that have bridged ANNs with neural data have primarily used architectures such as RNNs, MLPs, or CNNs^20,23,25,55–58^. The predominance of these models reflects their relative simplicity, as well as their historical roots in neuroscience^59–61^, whereas the transformer architecture was developed independently from neuroscience^13^.

Nevertheless, recent studies have identified potential links between transformers and specific neural systems, such as the hippocampus^31^, visual cortex^30^, cortico-thalamic circuits^29^, and language networks^62^. The relevance of transformers to relational reasoning lies in their ability to flexibly coordinate multiple individually ambiguous constraints over a 2D structural scaffold; a fundamental computational requirement of relational reasoning in humans. Critically, our results demonstrate these constraints can be understood in the context of the positional encoding and attention components of the transformer, offering a computational lens for probing analogous neurocognitive processes in the brain.

Neuroimaging work has primarily focused on identifying brain regions and networks that respond to increasing relational demands^7,8,40^, or that covary with individual differences in reasoning ability^63,64^, consistently highlighting the central role of a frontoparietal “multiple demand” network. We show that representations produced by the transformer’s attention mechanism align with representations generated by this network during relational reasoning. To understand how such structured representations could be generated computationally, we decomposed the transformer into its core components and examined their algorithmic contributions. Although the biological implementation is likely distinct, the transformer provides a useful high-level algorithmic analogue for how such representations could arise in the brain^65^.

Probing the attention mechanism demonstrated that increasing reasoning complexity was expressed as changes in information routing observed within the attention weights. Moreover, when attention weights were systematically manipulated (via the temperature parameter), more diffuse, higher entropy information routing degraded performance in such a way that mimicked human errors. Together, these results point towards a computational-level explanation for well-described capacity limits in human relational reasoning^1,34^.

In addition to the transformer’s attention mechanism, PE plays a critical role by grounding tokens in a stable spatial scaffold that attention can exploit. At an algorithmic level, PE can be viewed as operating analogously to spatial receptive fields^66^, providing a coordinate system that enables relational computations over structured inputs. This intuition is supported by recent work showing that transformers equipped with recurrent PEs can reproduce representational properties of the hippocampal formation during abstract spatial navigation tasks^31^. Here, our results demonstrate a related effect in a distinct domain: PE representations aligned with the grid structure of the stimulus space, and with representations observed in early visual cortex.

We also show that when forced to *learn* the correct 2D arrangement (using a learnable PE parameter), naive initialization does not guarantee strong model-brain alignment. Interpreted through recent theoretical work distinguishing lazy from rich learning in ANNs^50,51^, these results suggest sufficiently rich optimization dynamics are critical for PE to acquire task-relevant spatial structure. Together, these findings suggest that principles previously identified in feedforward networks—namely the importance of rich initializations for acquiring structured, brain-like representations^24^—extend to learnable PEs in transformers, which endow models with the requisite spatial representations necessary to perform 2D relational reasoning.

Our findings motivate several promising directions for future research. First, although the LST provides a tractable and well-studied model of relational reasoning, transformers have demonstrated the ability to learn even more complex reasoning tasks that have yet to be explored within neuroscience, such as Chess^67^, Go^68^, and the abstract reasoning corpus^69^.

Though designing feasible experiments to collect neural data for these tasks will be challenging, characterizing the neural mechanisms underlying reasoning across a diverse set of complex tasks that are both challenging for humans and artificial systems offers exciting directions for future work (for example increasing the LST puzzle size, as a form of size generalization^70,71^ will be challenging for both humans^72^ and models). Second, although standard implementations of MLPs and LSTMs failed to generalize on the LST, it is possible that non-standard training set ups, learning regimes, or architectural variants (e.g., graph^73^ or looped transformers^74^) may confer improved model performance or brain-model alignment. A systematic investigation of these possibilities could clarify what additional parameters are needed to enhance MLPs and LSTMs to natively perform the LST. Finally, though transformers were not designed with biological plausibility in mind, this does not preclude their utility as computational models for understanding neural systems and brain computation^19^, future work should explore more circuit-based, and biologically meaningful mechanisms through the use of biophysical models or, where possible, neural data collected from other animal models.

In summary, we investigated the computational mechanisms key for the relational reasoning required in the LST using artificial neural network models and fMRI data. Among commonly used neural architectures, we found that the transformer architecture enabled the most generalizable relational reasoning capability. Despite having origins outside of neuroscience, the transformer architecture offered a powerful computational framework for modeling relational reasoning and its neural correlates. By analyzing the contributions of the transformer’s core circuit mechanisms – PE and attention – to LST generalization, we revealed how these components jointly support the representation and manipulation of structured information, paralleling visual and cognitive control systems in the human brain. Moreover, the hierarchical progression of transformer representations across layers mirrored cortical processing hierarchies. This suggests shared algorithmic strategies in performing the LST in biological and gradient-descent-optimized artificial systems, underscoring the potential of transformer models as tools for probing the computational mechanisms of complex cognition. We hope this study inspires further work in leveraging more powerful artificial models to investigate the neural correlates of more sophisticated cognitive tasks.

## Methods

### Participants

Sixty-five healthy, right-handed adult participants completed the LST while undergoing fMRI. Of these, 40 were included in the final analysis (mean age=23 years old; age range=18 - 33; 60% female). Four participants were excluded due to MR scanning errors, one was excluded due to an unexpected brain structure abnormality, another was excluded for low accuracy in the behavioral task (total score more than three standard deviations below average) and 19 participants were excluded due to excessive head movement (note a stringent head motion threshold was employed - described below in the Neuroimaging section). Participants provided written informed consent and were eligible if they were aged between 18 - 35, with no previous reported history of mental health or neurological disorders. Ethics approval was granted by the University of Queensland Human Research Ethics Committee. This dataset has been used in previous publications^9,11,75^.

### Latin Square Task (LST) experiment design

The LST is a nonverbal relational reasoning task^32^. Each puzzle is a two-dimensional 4×4 grid populated with shapes (squares, triangles, circles, and crosses) and a target question mark. Participants must determine the shape of the target, ensuring each shape appears only once in every row and column, similar to Sudoku. Puzzles were organized into three conditions with increasing reasoning complexity^1^:

- One-vector: linking puzzle elements (shapes) across a single row or column
- Two-vector: linking elements across both a row and a column
- Three-vector: linking elements across three rows or columns.

Details of the fMRI implementation of the LST have been described previously^9,36^. In brief, 108 LST items (36 per relational complexity condition) were presented in the MR session across three runs. Participants viewed each puzzle for five seconds, followed by a brief pause and then a two second response screen. Motor responses were counterbalanced across participants, ensuring that an equal number of participants had the same shape-response mapping. Stimuli were presented using PsychToolBox 3.0^76^. In addition to the LST, participants also completed the Raven’s Advanced Progressive Matrices with a 40 minute time limit as a standard measure of fluid intelligence^77^.

### Neuroimaging

#### Acquisition

High resolution fMRI data were acquired on a 7 Tesla Siemens scanner equipped with a 32-channel head coil at the Centre of Advanced Imaging, The University of Queensland, Brisbane. Whole-brain echo-planar images were collected with the following parameters: isotropic 2 mm voxels, TR=586ms, TE= 23ms, flip angle=40°, FOV=208mm, 55 axial slices, multiband acceleration factor=5. Structural scans (MP2RAGE) used in the preprocessing pipeline were acquired with the following parameters: isotropic 0.75 mm voxels, TR=4300ms, TE=3.44 ms, and 256 slices.

#### Minimal preprocessing and denoising

Neuroimaging data were minimally preprocessed using a standard fMRIPrep pipeline (version 21.0.1)^78^ ^11^. In brief, this pipeline consisted of skull stripping, susceptibility distortion correction, coregistration, slice time correction and spatial normalisation (full details in **Supplementary Material**). The analyses in this manuscript focus on multi-vertex patterns, therefore any participants exceeding 2 mm of head motion in the six rotation and translation parameters were excluded from further analysis (N_excluded_=19, as in^11^). Additional denoising was performed by regressing fourteen nuisance signals (mean cerebrospinal fluid and white matter signals, along with the six motion parameters and their derivatives). Denoising was performed using the *clean* function in Nilearn^79^.

#### fMRI task activation estimation

Individual-level brain activations were estimated by conducting vertex-wise Least-Squares Separation (LSS)^80^ general linear model (GLM) analysis in grayordinate space^81^. A separate GLM was conducted for each puzzle in the experiment (N=108). Each design matrix included two regressors; one for the puzzle of interest, and one for all other puzzles in the experiment. This approach has been shown to improve the ability to detect single-trial activations over other standard methodologies^80^. Each puzzle was modeled as a five second boxcar regressor convolved with the Statistical Parametric Mapping^82^ canonical hemodynamic response function. Three additional regressors were included to account for each run of the experiment. Task GLMs were performed using the *run_glm* function in Nilearn^79^. In subsequent analysis of fMRI data we adopted the Glasser cortical parcellation^81^ by analyzing vertices within each parcel. We used the Cole–Anticevic network parcellation^47^ to contextualize our results within established functional brain networks.

### Artificial Neural Networks (ANNs)

We trained three ANNs with distinct architectures to perform the LST. Three thousand training LST puzzles were generated for each of the one-, two-, or three-vector conditions. The aforementioned 108 fMRI experiment puzzles were used for validation and comparison to empirical data. We ensured that the similarity between the training and validation puzzles was low (Jaccard dissimilarity > 0.8). For all models, we used the Adam optimizer with a learning rate of 0.0001 without regularization (no dropout or weight decay). We trained on 15 independent seeds for a fixed number of epochs (4000). Each model trained on 8000 of the original 9000 puzzles, holding out 1000 puzzles as an additional test set.

#### Transformer

The input to the transformer was a one-hot encoding of the symbol for each element in the grid. Since there were five possible symbols, each token was represented as a 5-dimensional vector.

If the grid element was empty, the token was a vector of zeros. We used a standard encoder-only transformer architecture with four layers and model embedding dimension of 160 ^13^. The context window for the model was 16 tokens long, given the two-dimensional 4 × 4 structure of the LST. Each model contained fully-connected bidirectional attention. For simplicity (and interpretability) we limited the number of attention heads to 1. We used the standard position-wise MLP in the transformer with a single hidden layer with an expansive embedding dimension that was four times larger than the token embedding dimension (160*4) and a GELU nonlinearity. For PE, we used a 2D absolute PE and a separate learnable embedding (described in more detail below). Both these PE variants were incorporated to the token embedding via vector addition.

More formally, let *X* be a 16 × 160 matrix (number of tokens × embedding dimensions) that denotes our input tokens, *P* be our PE matrix of the same dimensions, **A** be the attention mechanism (defined here as a function), and **M** be the MLP (defined as a function). Then, a single transformer layer is computed as

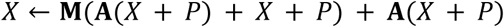

where the additional *X* + *P* and **A**(*X* + *P*) terms are standard residual connections that are common in transformers. We define the attention mechanism **A** as 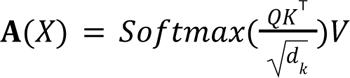, where *Q* = *XW_Q_*, *K* = *XW_K_*, and *V* = *XW_V_* with *W_Q_*, *W_K_*, *W_K_* as learnable *d* × *d* (i.e., 160 x 160) matrices. The Softmax is a row-wise computation (applied to row vectors) defined as 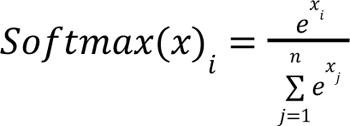, where *x* is a row vector in the *n* × *n* attention matrix (*n* = 16 for the LST task), and yields probability values (summed to 1) for all elements in the row vector. The intuition is that the highest probability value is assigned to the token that is most strongly attended to.

The function **M** is a learnable MLP specified above with a single hidden layer. Note that we assume LayerNorms, a common normalization technique in transformers, to be carried out after functions **A** and **M**. We use standard placements of the LayerNorms (as specified in Vaswani et al., 2017), which can be found in the code.

#### Positional encoding (PE) in transformers

In this analysis we explored the impact of PE on downstream model performance and representations. The two-dimensional LST puzzle grid can be explicitly modeled in the PE of the transformer. In this context, the PE reflects the model’s ability to grasp the 2D task structure; without an understanding of the puzzle’s square layout, it cannot access the row and column information needed to solve it visually. While most implementations of PEs employed by transformers are optimized for 1D text^13,83–85^, a number of studies have demonstrated the importance of PE for more structured tasks (rather than natural language), such as arithmetic^86–88^, or tasks requiring multiple dimensions (e.g., 2D tasks). Two-dimensional absolute PEs are a generalization of 1d-fixed presented in prior work^13^. The primary distinction is that, rather than using a single position variable where 1<*pos*< 16, there are two position variables: 1≤*pos_w_*≤ 4 and 1≤*pos_h_*≤ 4, representing the width and height of the grid, respectively. Positional encoding for a row *w* is defined by:

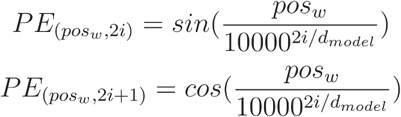

Positional encoding for a column *h* is defined by

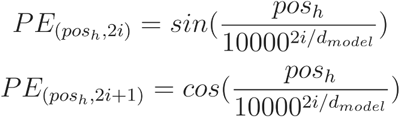

For a 2D PE, half of the embedding dimensionality is allocated to encoding rows, and the other half to columns. Thus, for *d_model_* =160, embedding dimensions 0–79 are reserved for row encoding and embedding dimensions 80–159 are reserved for column encoding. This PE formulation provides a strong inductive bias for 2D reasoning problems.

This inductive bias may make comparisons with our control ANNs less direct, as those models do not explicitly encode 2D structure. Unlike transformers with a 2D PE, MLPs and LSTMs need to learn a 2D representation via training. Nevertheless, prior work shows that analogous constraint-satisfaction problems like Sudoku can be formulated as 1D strings^42^, and that state-of-the-art symbolic solvers operate solely on 1D inputs without explicit positional structure^89^. To control for this potential bias, we used a transformer with no positional inductive bias, and instead a learnable free PE parameter (represented as a 160-dimensional vector per token). Prior work in deep learning theory suggests that the choice of weight initialization can influence both the structure of learned representations and task generalization ^50,51,90^.

Specifically, smaller initialization norms tend to promote more structured internal representations. We assessed whether these findings would extend to learnable transformer PEs initialized from different Normal distributions, controlling for the standard deviation (i.e., norm). For each token embedding, we initialized a learnable PE parameter from a multivariate Normal distribution, denoted *N*(0,Σ), where Σ=σI, and I denotes the identity matrix scaled by σ As in the prior analysis, we trained 15 independent seeds across 4000 epochs. All models converged in training. We then tested whether there was an interaction between reasoning complexity condition and initialization norm in regards to model generalization accuracy. This PE analysis was specific to the results presented alongside **Fig. 3**, whereas the fixed 2D PE was used for all other analyses.

#### Control ANNs

Two control ANNs were assessed in their ability to complete the LST, i) a feedforward fully-connected MLP and, ii) a bidirectional Long Short-Term Memory (LSTM) network. Each of the control models contained an input layer, four hidden layers of 160 units, and an output layer. Both ANNs were implemented with ‘out-of-the-box’ PyTorch models, and are included with the code ^91^. Despite all models and seeds converging during training, these models did not exhibit meaningful generalization to unseen LST puzzles.

The input to the MLP was a concatenation of the one-hot encodings of all grid elements. As there were 5 possible symbols (per grid element) and 16 possible grid locations, this yielded an 80-dimensional input vector. The input to the LSTM (as a sequence model) was to present each LST grid element (as a 5-dimensional vector) in sequence, both forwards and backwards (i.e., bidirectionally). In other words, the LSTM recurrently iterated 32 (16*2) times prior to forming a decision. This contrasts with the transformer, where each token is processed separately (but in parallel) and PE is applied separately to each token. In the MLP and LSTM, grid position information is implicitly encoded, as each token has its own set of unique input channels/weights in the MLP, and each token is processed in ordered sequence in the LSTM. Thus, MLPs and LSTMs are conceptually similar to transformers with a learnable PE, as those models need to learn which set of input channels/tokens belong to which grid position implicitly via weight updates. Note, however, that the learnable PE in transformers is a vector parameter, whereas in MLPs and LSTMs it learns grid position through weight updates on the input embeddings. In the absence of the ability to accurately learn the correct 2D positional structure, the formulation of the MLP effectively encodes a randomized PE of grid elements, as all input channels are orthogonal to each other at initialization.

### Representational Similarity Analysis (RSA)

We used RSA to contrast experimenter task models, transformer model activations and fMRI brain activations ^43,44^. All models and representational dissimilarity matrices (RDMs) were derived across the 108 test LST puzzles. Three task models were contrasted; stimulus, complexity and response (shown in **Fig. 2D**). The stimulus model was calculated with the *jaccard distance* between each of the 108 one-hot encoded puzzles. Specifically:

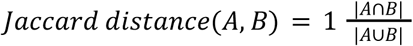

Thus, puzzles with similar layouts, regardless of their complexity, had scores closer to 0 and vice versa. The relational complexity model represented the different conditions of the experiment whereby each condition was coded to be maximally similar (0), adjacent conditions were dissimilar (0.5, e.g. one-vector vs. two vector) and one-vector and three-vector puzzles were maximally dissimilar (1). Finally, as in the stimulus model, the response model was calculated with the jaccard distance between each of the 108 one-hot encoded solutions. Thus, puzzles with the same target response (e.g., square) would be considered similar. As in other RSA studies, we ensured low-level task features (stimulus or response) were not associated with the relational complexity model. Specifically, relational complexity was not predicted by the raw stimulus input (R^2^_adj_=.296, F(80,28)=1.57, *p*=.09) or raw response output (R^2^_adj_=-.025, F(4,104)=0.13, *p*=.94). Likewise, at the representational level, neither the stimulus RDM (R^2^_adj_=-.0002, F(1,5776)=0.31, *p*=.86) nor the response RDM (R^2^_adj_≈0, F(1,5776)=0.95, *p*=.33) was associated with the relational complexity model.

For the transformer and fMRI data we used *rsatoolbox* to generate representations^44^. Specifically, given there is no noise in the measurement of transformer neural activity, we used the squared Euclidean distance to derive RDMs. This was performed for token embeddings after application of the PE (*X_PE_*= *e*(*x_i_*) + *p_i_*) and Attention mechanisms ( *X_Attn_* = *softmax*(*QK^T^*)*V* + *X_PE_*). For the fMRI data, participants did not see any puzzles more than once, to ensure they couldn’t trivially memorise puzzle geometries and solutions. Thus, cross-validation was performed across-participants (rather than *within-participant*) using the squared Mahalanobis distance (“crossnobis” distance). This resulted in a single RDM per brain region in the parcellation. The upper triangle of the RDMs were compared using Pearson correlation; we refer to this as “representational alignment”^92^.

#### Experimenter model comparisons

The primary analysis using the experimenter models was to evaluate which embeddings from each transformer component (PE, or Attention), and model layer (1 - 4) were most similar to the experimenter models (stimulus, complexity, response). To test this, alignment values (i.e., Pearson’s *r*) were submitted to a repeated-measures ANOVA (2 x component, 4 x layer), with model seeds treated as subjects. A control analysis was conducted wherein we evaluated the representational alignment between models that had undergone a single epoch of training and fully trained models (4000 epochs). We also contrasted the control MLP model with the Transformer. These results are shown in **Supplementary Fig. 1**, and **Tables 2-3**.

#### Brain-transformer comparisons

For the comparison between transformer and brain imaging, significance was ascribed using a permutation method, whereby the condition labels (i.e., puzzles) were reordered within the transformer RDMs, and then compared with the empirical fMRI RDM. This was repeated 10,000 times to generate a null distribution of unrelated RDMs for every brain region^93^. For a given brain region, if the true correlation exceeded the 95th percentile, the null hypothesis was rejected. In addition to this procedure, we performed false discovery rate (FDR) correction across the 360 brain regions in the brain parcellation.

For the main analysis, the PE and Attention embeddings were averaged across layers prior to RDM calculation. For the later, layer-dependent analysis (described in “Brain-transformer alignment across layers and comparison to cortical gradients”), this averaging step was not performed. We also examined the relationship between brain-transformer (PE embeddings) representational alignment and initialization norms within regions of interest which had shown an effect in the main analysis (i.e., the primary visual cortex).

A control analysis was performed to ensure stimulus, complexity and response information were detectable in the fMRI data. Specifically, decoding was performed for each participant using logistic regression in a leave-one-puzzle-out cross-validation scheme. Classification accuracy was averaged across folds and compared against chance using one-sample *t*-tests. Chance levels for relational complexity and motor response decoding were 33%(1-, 2-, 3-vector) and 25% (four possible button responses), respectively. For the stimulus model, because each puzzle was presented only once per participant, one-hot encoded puzzle features were clustered using k-means, yielding 38 clusters based on the highest silhouette score and a corresponding decoding chance level of 2.84% for visual decoding. Decoding was assessed separately for each cortical parcel (n = 360), with significance determined using FDR correction across regions. (**Supplementary Fig. 5 and Supplementary Table 7**).

A further control analysis was conducted wherein we evaluated the representational alignment between the fMRI RDMS and untrained transformer RDMs (single epoch of training). This analysis followed the same procedure as above. These results are shown in **Supplementary Fig. 2**.

#### Noise ceiling estimation

Noise ceilings aim to estimate the degree to which a theoretically ideal model, subject to the same data and measurement constraints, could explain the observed representational structure. For non-fMRI analyses (e.g., **Fig. 2E**), the theoretical noise ceiling is 1. For example, if a model performed no transformation in its first layer and simply passed the input through unchanged, its representations would be identical to those of the stimulus model (**Fig. 2D**), yielding perfect representational alignment. For the fMRI analyses, we estimated noise ceilings using a split-half reliability approach, similar to prior work^94^. Specifically, for each brain region, participant data were randomly split into two independent groups (n = 20 per split), and crossnobis RDMs were computed separately for each group. The upper triangles of the two RDMs were then correlated, and the resulting correlation coefficient was adjusted for reduced sample size using the Spearman–Brown correction. This procedure was repeated 1000 times for each region to generate a distribution of noise-ceiling estimates.

### Attention weight analysis

In this analysis we sought to relate attention weights (*QK^T^*) to relational complexity in the LST. Attention weights are the token by token (in our case 16 x 16) matrices denoting how each puzzle element relates to one another. These values were averaged over layers. We analyzed the mean *key* attention (MKA) weights (shown in **Fig. 4A**), which represent the extent to which each token broadcasts their information to all other tokens. The focus on keys was important for two considerations. Conceptually, information broadcasting naturally aligns with the cognitive integration demands of relational complexity. **Technically**, we emphasize keys rather than queries because of the properties of the softmax operation applied to attention weights. In standard attention mechanisms, the softmax is applied independently for each query, enforcing a row-wise normalization such that attention weights sum to one. This constraint makes differences in attention along the query dimension difficult to interpret in the present context.

We performed three complementary analyses. First, we conducted a repeated measures *t*-test comparing MKA weights for the target (i.e. “?”) and non-target tokens. Second, we examined whether the cells critical for solving a puzzle had higher attention weights than non-critical cells. Critical cells were defined as elements in the grid that directly contributed to the prediction of the element in the target location. We only evaluated one-vector puzzles where the critical cells are tractable; they will always occur within the same row or column as the target. We conducted a repeated measures *t*-test to compare the MKA weights of critical and non-critical cells. Lastly, we tested whether the non-target MKA weights differed across the three complexity conditions via a Friedman test, with follow-up *t*-tests between conditions.

As a final analysis of the attention weights, we manipulated the softmax’s *temperature* parameter as a mechanistic probe of how “attention spread” relates to task performance and its correspondence with human behavior. Specifically, the *n* by *n* attention weights *A* are calculated as:

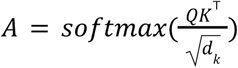

where an element *a* in *A* is calculated as:

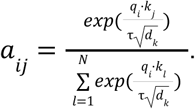

When temperature τ = 1 (the default parameter used during learning), attention weights were unaltered. Lower temperatures (τ < 1) sharpen the resulting attention distribution by concentrating probability mass on a small subset of tokens, whereas higher temperatures ( τ > 1) produce progressively flatter, higher-entropy attention distributions. Manipulating the temperature of the model did not require retraining. We evaluated a wide range of temperatures (τ = 0. 0001 to τ = 1000) to characterize the full regime of model behavior (**Fig. 4E**). Across temperatures, we assessed model accuracy on held-out puzzles and additionally examined model–human correspondence by correlating Transformer accuracy (averaged across random seeds) with human accuracy (averaged across participants) on a per-puzzle basis (**Supplementary Fig. 4** and **Supplementary Table 6)**.

### Brain-transformer alignment across layers and comparison to cortical gradients

In the final analysis we tested brain-transformer alignment across each of the model’s layers, and compared the corresponding brain maps to connectivity-based cortical gradients. In this analysis, we examined embeddings from the final MLP architectural component of the transformer and estimated representational alignment to fMRI data as described previously. We then correlated these brain surface maps with the first two connectivity gradients derived from prior work^53^ using neuromaps^54^. Finally, statistical significance was ascribed by generating a null distribution from 10,000 spin-test permutations^95^, in which the cortical gradient maps were randomly rotated on the surface and correlated with the ANN–fMRI alignment maps, with significance subsequently corrected for eight multiple comparisons (two gradients by four layers) using FDR.

## Data availability statement

Raw human behavioral and fMRI data is available online via The University of Queensland’s espace library (http://dx.doi.org/10.14264/uql.2019.780). Training and testing data for the neural network models are available via the repository on github (https://github.com/ljhearne/LSTNN_public).

## Code availability statement

Code used to perform the analyses are available via the repository on github (https://github.com/ljhearne/LSTNN_public).

## Acknowledgements

This work was supported by the Australian NHMRC (LC: GN2001283 and GNT2027597, LJH: APP1194070).

## Author contributions

LJH and TI conceptualized and designed the project. LJH and LC collected the data. LJH, CR, and TI analysed the data. LJH and TI wrote the original draft. All authors edited the manuscript.

