## Supplemental Information for "Aligning transformer circuit mechanisms to neural representations in relational reasoning"

**Transformer model performance**
Given the highly non-normal distribution of accuracy values, we used the non-parametric Friedman test. There was an effect of reasoning complexity on Transformer accuracy ($\chi^2(2)=11.47$, $p=.003$). Follow up *t*-tests revealed a difference between 2-vector (*M*=97.78%, *SD*=8.31%) and 3-vector puzzles (*M*=96.11%, *SD*=9.56%, $t(14)=2.81$, $p=.014$, $d=0.18$) but not 1-vector (*M*=99.07%, *SD*=3.46%) and 2-vector puzzles ($t(14)=1.0$, $p=.334$, $d=0.20$), possibly due to ceiling effects on accuracy. While this provides some evidence for a performance cost due to relational complexity, the number of errors in the more difficult condition was much smaller than in humans.

**Representational similarity control analyses**
In addition to the results presented in the main text, we compared the experimenter-designed models with an untrained transformer (trained for a single epoch). We also contrasted the final trained transformer output (MLP block) and a control MLP model. These analyses are presented in **Supplementary Fig. 1 and Supplementary Tables 2–3**. We also contrasted untrained transformer models with fMRI representations using the same analysis pipeline as in the main manuscript (**Supplementary Fig. 2**).

These analyses show that untrained and poorly performing models (i.e., the control MLP) can still capture features of the input stimulus. Subsequently, when contrasted with fMRI data, the untrained transformer can capture stimulus-related variance in the fMRI data (i.e., significant representational alignment in right V1). This result is not unexpected, given that poor performing models likely pass the stimulus input through the model without learning the appropriate weight adjustments to solve the LST (e.g., the untrained attention embedding RSA show a similar pattern to that of the untrained PE embedding RSA in the figure above, as nothing has been learned). The analyses also demonstrate that successful training increased the alignment between the transformer and the stimulus model, as well as fMRI data in the visual cortex. Moreover, training is necessary for the transformer embeddings to capture puzzle complexity and response models, and to align attention embeddings with fMRI data outside of the visual cortex.

**fMRI preprocessing**

Results included in this manuscript come from preprocessing performed using fMRIPrep 21.0.1 (Esteban, Markiewicz, et al. (2018); Esteban, Blair, et al. (2018); RRID:SCR_016216), which is based on Nipype 1.6.1 (K. Gorgolewski et al. (2011); K. J. Gorgolewski et al. (2018); RRID:SCR_002502). Before running fMRIPrep, the raw T1-weighted (T1w) images were intensity-corrected using the SPM segmentation tool to address signal intensity differences between the temporal cortex and the rest of the brain. Quality control checks with and without this step confirmed improved tissue classification accuracy during the subsequent preprocessing pipeline.

*Anatomical data preprocessing*

A total of 1 T1-weighted (T1w) images were found within the input BIDS dataset.The T1-weighted (T1w) image was corrected for intensity non-uniformity (INU) with N4BiasFieldCorrection (Tustison et al. 2010), distributed with ANTs 2.3.3 (Avants et al. 2008, RRID:SCR_004757), and used as T1w-reference throughout the workflow. The T1w-reference was then skull-stripped with a Nipype implementation of the antsBrainExtraction.sh workflow (from ANTs), using OASIS30ANTs as target template. Brain tissue segmentation of cerebrospinal fluid (CSF), white-matter (WM) and gray-matter (GM) was performed on the brain-extracted T1w using fast (FSL 6.0.5.1:57b01774, RRID:SCR_002823, Zhang, Brady, and Smith 2001). Brain surfaces were reconstructed using recon-all (FreeSurfer 6.0.1, RRID:SCR_001847, Dale, Fischl, and Sereno 1999), and the brain mask estimated previously was refined with a custom variation of the method to reconcile ANTs-derived and FreeSurfer-derived segmentations of the cortical gray-matter of Mindboggle (RRID:SCR_002438, Klein et al. 2017). Volume-based spatial normalization to two standard spaces (MNI152NLin2009cAsym, MNI152NLin6Asym) was performed through nonlinear registration with antsRegistration (ANTs 2.3.3), using brain-extracted versions of both T1w reference and the T1w template. The following templates were selected for spatial normalization: ICBM 152 Nonlinear Asymmetrical template version 2009c [Fonov et al. (2009), RRID:SCR_008796; TemplateFlow ID: MNI152NLin2009cAsym], FSL's MNI ICBM 152 non-linear 6th Generation Asymmetric Average Brain Stereotaxic Registration Model [Evans et al. (2012), RRID:SCR_002823; TemplateFlow ID: MNI152NLin6Asym].

*Functional data preprocessing*

For each of the 3 BOLD runs found per subject (across all tasks and sessions), the following preprocessing was performed. First, a reference volume and its skull-stripped version were generated using a custom methodology of fMRIPrep. Head-motion parameters with respect to the BOLD reference (transformation matrices, and six corresponding rotation and translation parameters) are estimated before any spatiotemporal filtering using mcflirt (FSL 6.0.5.1:57b01774, Jenkinson et al. 2002). BOLD runs were slice-time corrected to 0.268s (0.5 of slice acquisition range 0s-0.535s) using 3dTshift from AFNI (Cox and Hyde 1997, RRID:SCR_005927). The BOLD time-series (including slice-timing correction when applied) were resampled onto their original, native space by applying the transforms to correct for head-motion. These resampled BOLD time-series will be referred to as preprocessed BOLD in original space, or just preprocessed BOLD. The BOLD reference was then co-registered to the T1w reference using bbregister (FreeSurfer) which implements boundary-based registration (Greve and Fischl 2009). Co-registration was configured with six degrees of freedom. Several confounding time-series were calculated based on the preprocessed BOLD: framewise displacement (FD), DVARS and three region-wise global signals. FD was computed using two formulations following Power (absolute sum of relative motions, Power et al. (2014)) and Jenkinson (relative root mean square displacement between affines, Jenkinson et al. (2002)). FD and DVARS are calculated for each functional run, both using their implementations in Nipype (following the definitions by Power et al. 2014). The three global signals are extracted within the CSF, the WM, and the whole-brain masks. Additionally, a set of physiological regressors were extracted to allow for component-based noise correction (CompCor, Behzadi et al. 2007). Principal components are estimated after high-pass filtering the preprocessed BOLD time-series (using a discrete cosine filter with 128s cut-off) for the two CompCor variants: temporal (tCompCor) and anatomical (aCompCor). tCompCor components are then calculated from the top 2% variable voxels within the brain mask. For aCompCor, three probabilistic masks (CSF, WM and combined CSF+WM) are generated in anatomical space. The implementation differs from that of Behzadi et al. in that instead of eroding the masks by 2 pixels on BOLD space, the aCompCor masks are subtracted a mask of pixels that likely contain a volume fraction of GM. This mask is obtained by dilating a GM mask extracted from the FreeSurfer's aseg segmentation, and it ensures components are not extracted from voxels containing a minimal fraction of GM. Finally, these masks are resampled into BOLD space and binarized by thresholding at 0.99 (as in the original implementation). Components are also calculated separately within the WM and CSF masks. For each CompCor decomposition, the k components with the largest singular values are retained, such that the retained components' time series are sufficient to explain 50 percent of variance across the nuisance mask (CSF, WM, combined, or temporal). The remaining components are dropped from consideration. The head-motion estimates calculated in the correction step were also placed within the corresponding confounds file. The confound time series derived from head motion estimates and global signals were expanded with the inclusion of temporal derivatives and quadratic terms for each (Satterthwaite et al. 2013). Frames that exceeded a threshold of 0.5 mm FD or 1.5 standardised DVARS were annotated as motion outliers. The BOLD time-series were resampled into standard space, generating a preprocessed BOLD run in MNI152NLin2009cAsym space. The BOLD time-series were resampled onto the following surfaces (FreeSurfer reconstruction nomenclature): fsaverage. Grayordinates files (Glasser et al. 2013) containing 91k samples were also generated using the highest-resolution fsaverage as intermediate standardized surface space. All resamplings can be performed with a single interpolation step by composing all the pertinent transformations (i.e. head-motion transform matrices, susceptibility distortion correction when available, and co-registrations to anatomical and output spaces). Gridded (volumetric) resamplings were performed using antsApplyTransforms (ANTs), configured with Lanczos interpolation to minimize the smoothing effects of other kernels (Lanczos 1964). Non-gridded (surface) resamplings were performed using mri_vol2surf (FreeSurfer).

Many internal operations of fMRIPrep use Nilearn 0.8.1 (Abraham et al. 2014, RRID:SCR_001362), mostly within the functional processing workflow. For more details of the pipeline, see the section corresponding to workflows in fMRIPrep's documentation.

Copyright Waiver

The above boilerplate text was automatically generated by fMRIPrep with the express intention that users should copy and paste this text into their manuscripts unchanged. It is released under the CC0 license.

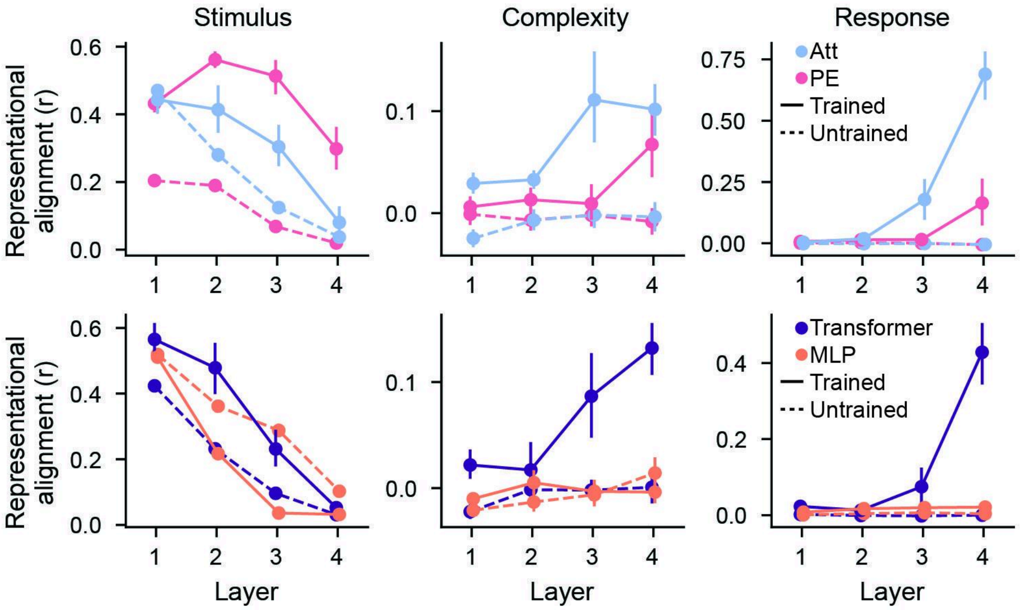


**Supplementary Fig. 1.** *Representational alignment between task feature models and baseline models.* The top row shows embedding alignment for trained (solid line; result from **Fig. 2E**) and untrained models (dashed line) Transformer components (PE; pink, attention; blue). The bottom row shows the comparison between trained and untrained Transformer (purple) and MLP models (orange). Error bars show bootstrapped 95% confidence intervals.

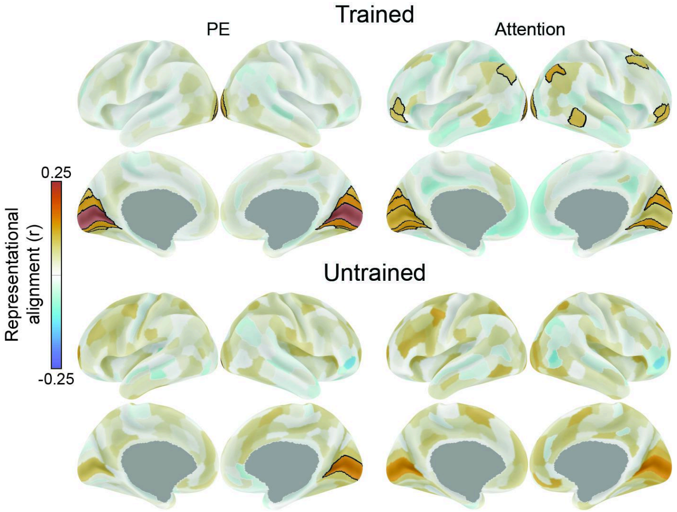


**Supplementary Fig. 2.** *Representational alignment between brain and transformer (PE and Attention) embeddings for trained (top panel) and untrained models (bottom panel).* The top panel shows the same analysis as **Fig. 2G** for reference. The bottom panel shows the alignment between untrained transformer models and fMRI data. The methodology is otherwise identical to the top panel. Black borders indicate significant ANN-fMRI representational alignment (permutation and FDR corrected). A single brain region contrasting PE embeddings and the untrained transformer reached statistical significance; right V1, $r$=.17, $p_{FDR}$<.001. For reference, the same contrast in the trained model; right V1, r=.26, $p_{FDR}$ <.001.

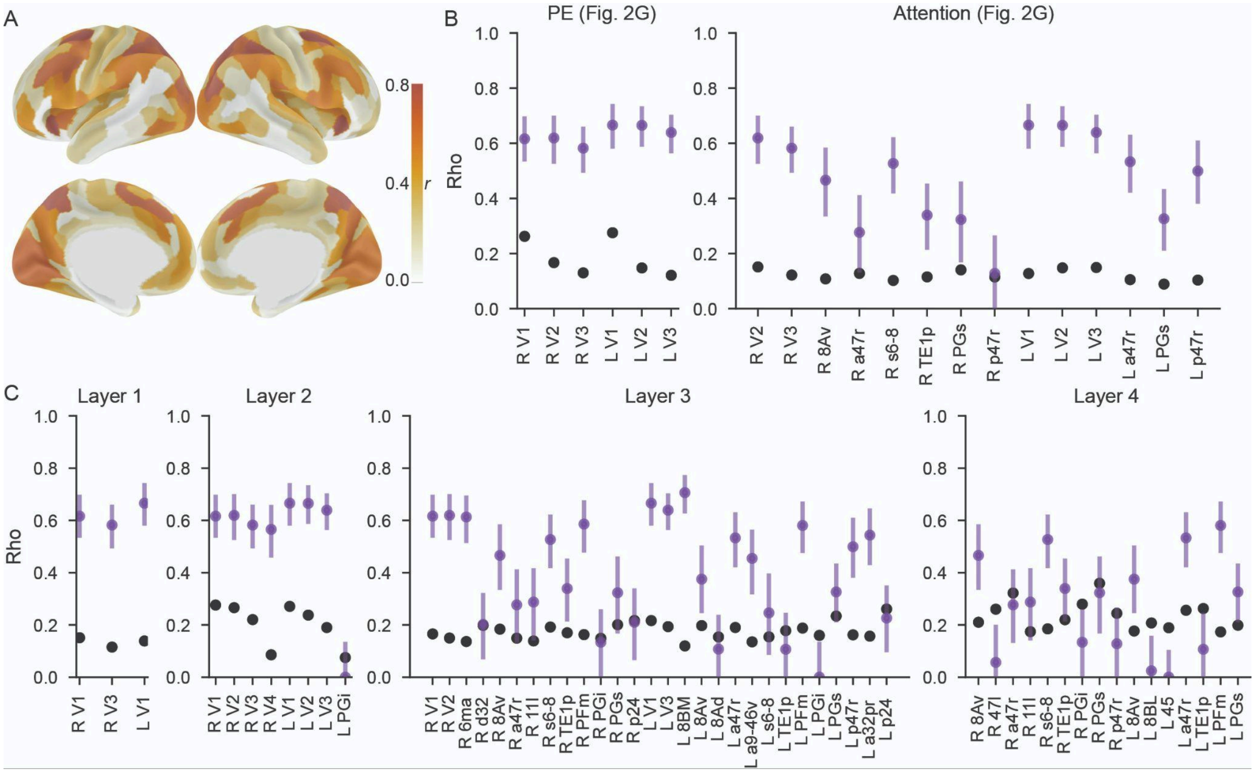


**Supplementary Fig. 3.** *Split-half noise ceiling estimation in brain imaging data.* **A.** Average noise ceiling (spearman-brown corrected) values across 1000 permutations for all cortical regions. **B.** Individual region noise ceiling estimates for significant brain regions in **Fig. 2G. C.** Individual region noise ceiling estimates for significant brain regions in **Fig. 5.** Black data show the empirical results. Purple data indicates the noise ceiling estimate. Error bars represent the 5th and 95th percentile across the 1000 permutations.

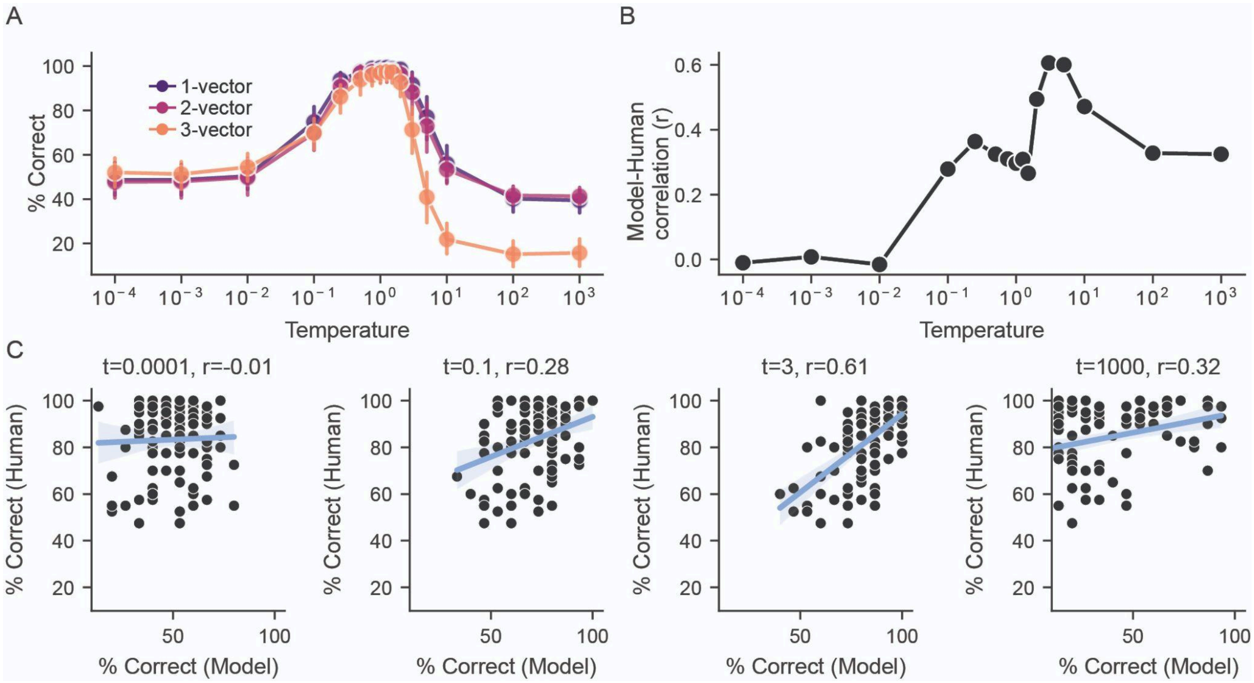


**Supplementary Fig. 4.** *Impact of temperature on model performance and behavioural associations.* **A.** Trained transformer model performance when the temperature hyperparameter is manipulated. Good model accuracy is achieved near $\tau = 1$. Both high and low temperatures caused decreased performance. **B.** Correlation between averaged human accuracy (N=40) and averaged transformer accuracy (N=15) across temperature manipulations. Peak correlation was observed in higher temperatures ($\tau = 3$). **C.** Example correlations from panel **B**.

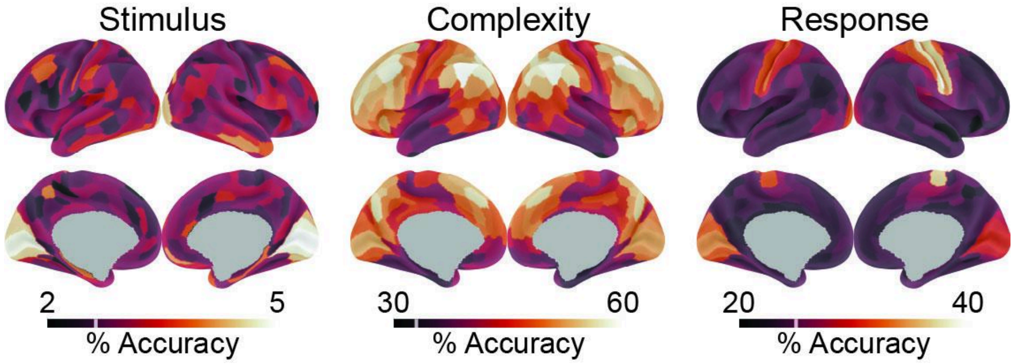


**Supplementary Fig. 5.** Decoding stimulus, complexity and response categories from fMRI data. A leave-one-puzzle-out decoder was trained for each participant to predict either stimulus (left), relational complexity (middle) or response categories (right). Chance accuracy was 2.84%, 33%, and 25% for each of the analyses. Lighter colours indicate higher decoding performance. As expected, peak decoding of the stimulus and motor response information was in the primary visual cortex (mean accuracy=4.75%, R_V1, $p_{FDR} < 0.001$) and the primary sensory cortex (mean accuracy=37.58%, R_3b, $p_{FDR} < 0.001$), respectively. Reasoning complexity decoding was observed in largely frontoparietal brain areas, with peak decoding in the lateral prefrontal cortex (58.43%, R_8Av, $p_{FDR} < 0.001$).

**Supplementary Table 1.** Representational alignment between transformer components and experimenter models (stimulus, complexity, response)

| **Task Feature** | **Source** | **SS** | **df₁** | **df₂** | **MS** | **F** | ***p*** | $\eta^2_G$ |
|---|---|---|---|---|---|---|---|---|
| Visual | Layer | 1.563 | 3 | 42 | 0.521 | 76.969 | < 0.001 | 0.611 |
| | Mechanism | 0.594 | 1 | 14 | 0.594 | 575.267 | < 0.001 | 0.374 |
| | Interaction | 0.253 | 3 | 42 | 0.084 | 9.317 | < 0.001 | 0.203 |
| Complexity | Layer | 0.091 | 3 | 42 | 0.03 | 15.268 | < 0.001 | 0.279 |
| | Mechanism | 0.06 | 1 | 14 | 0.06 | 131.594 | < 0.001 | 0.204 |
| | Interaction | 0.034 | 3 | 42 | 0.011 | 5.038 | 0.022 | 0.125 |
| Response | Layer | 3.524 | 3 | 42 | 1.175 | 103.725 | < 0.001 | 0.711 |
| | Mechanism | 0.894 | 1 | 14 | 0.894 | 144.919 | < 0.001 | 0.384 |
| | Interaction | 1.376 | 3 | 42 | 0.459 | 37.32 | < 0.001 | 0.49 |

Note: As explained in the main text, each task feature was analysed with a 4 (layer) by 2 (mechanism; PE vs. attention) repeated-measures ANOVA. Fifteen seeds were treated as subjects. SS = Sum of Squares; $df_1$, $df_2$ = degrees of freedom; MS = Mean Square;p = Greenhouse–Geisser corrected p-value; $\eta^2 g$ = generalized eta squared effect size.

**Supplementary Table 2.** Average representational alignment values between trained and untrained transformer components and experimenter models (stimulus, complexity, response)

| **Experimenter model** | **Component** | **Training** | **Mean** | **STD** | **SEM** |
|---|---|---|---|---|---|
| Stimulus | Attention | Untrained | 0.228 | 0.168 | 0.022 |
| | | Trained | 0.31 | 0.175 | 0.023 |
| | PE | Untrained | 0.12 | 0.082 | 0.011 |
| | | Trained | 0.451 | 0.131 | 0.017 |
| Complexity | Attention | Untrained | -0.009 | 0.023 | 0.003 |
| | | Trained | 0.069 | 0.064 | 0.008 |
| | PE | Untrained | -0.004 | 0.021 | 0.003 |
| | | Trained | 0.024 | 0.045 | 0.006 |
| Response | Attention | Untrained | -0.002 | 0.008 | 0.001 |
| | | Trained | 0.223 | 0.308 | 0.04 |

| | | | | | |
|---|---|---|---|---|---|
| | PE | Untrained | 0 | 0.012 | 0.002 |
| | | Trained | 0.05 | 0.112 | 0.014 |

**Supplementary Table 3.** Average representational alignment values between trained and untrained transformer and multilayer perception and experimenter models (stimulus, complexity, response)

| Experimenter model | ANN | Training | Mean | STD | SEM |
|---|---|---|---|---|---|
| Stimulus | MLP | Untrained | 0.318 | 0.153 | 0.02 |
| | MLP | Trained | 0.199 | 0.198 | 0.026 |
| | Transformer | Untrained | 0.196 | 0.153 | 0.02 |
| | Transformer | Trained | 0.332 | 0.228 | 0.029 |
| Complexity | MLP | Untrained | -0.006 | 0.023 | 0.003 |
| | MLP | Trained | -0.003 | 0.018 | 0.002 |
| | Transformer | Untrained | -0.006 | 0.022 | 0.003 |
| | Transformer | Trained | 0.064 | 0.071 | 0.009 |
| Response | MLP | Untrained | 0.004 | 0.01 | 0.001 |
| | MLP | Trained | 0.016 | 0.013 | 0.002 |
| | Transformer | Untrained | 0 | 0.009 | 0.001 |
| | Transformer | Trained | 0.135 | 0.194 | 0.025 |

**Supplementary Table 4.** Significant representational alignment between transformer mechanisms and cortical regions

| Mechanism | Region | Network | Representational alignment (r) | $p_{FDR}$ | Noise Ceiling Mean (5th -95th percentile) | Percentage of upper Noise Ceiling |
|---|---|---|---|---|---|---|
| Attention | R_8Av | Default | 0.108 | < 0.001 | 0.47 (0.34-0.58) | 18.72 |
| Attention | R_PGs | Default | 0.141 | < 0.001 | 0.32 (0.18-0.45) | 31.07 |
| Attention | L_PGs | Default | 0.089 | 0.0499 | 0.33 (0.22-0.43) | 20.86 |
| Attention | R_a47r | Frontoparietal | 0.128 | 0.0212 | 0.28 (0.14-0.4) | 31.66 |
| Attention | R_s6-8 | Frontoparietal | 0.102 | 0.0212 | 0.53 (0.43-0.61) | 16.61 |
| Attention | R_TE1p | Frontoparietal | 0.115 | 0.0388 | 0.34 (0.22-0.45) | 25.79 |
| Attention | R_p47r | Frontoparietal | 0.115 | < 0.001 | 0.13 (-0.0-0.26) | 44.57 |

| | | | | | | |
|---|---|---|---|---|---|---|
| Attention | L_a47r | Frontoparietal | 0.105 | < 0.001 | 0.53 (0.43-0.62) | 16.85 |
| Attention | L_p47r | Frontoparietal | 0.104 | 0.0499 | 0.5 (0.39-0.6) | 17.27 |
| Attention | L_V1 | Visual1 | 0.128 | < 0.001 | 0.67 (0.59-0.73) | 17.42 |
| Attention | R_V2 | Visual2 | 0.151 | < 0.001 | 0.62 (0.53-0.69) | 21.81 |
| Attention | R_V3 | Visual2 | 0.122 | < 0.001 | 0.58 (0.5-0.65) | 18.71 |
| Attention | L_V2 | Visual2 | 0.148 | < 0.001 | 0.67 (0.6-0.73) | 20.36 |
| Attention | L_V3 | Visual2 | 0.149 | < 0.001 | 0.64 (0.57-0.7) | 21.42 |
| PE | R_V1 | Visual1 | 0.262 | < 0.001 | 0.62 (0.54-0.69) | 37.96 |
| PE | L_V1 | Visual1 | 0.276 | < 0.001 | 0.67 (0.59-0.73) | 37.56 |
| PE | R_V2 | Visual2 | 0.167 | < 0.001 | 0.62 (0.53-0.69) | 24.12 |
| PE | R_V3 | Visual2 | 0.13 | < 0.001 | 0.58 (0.5-0.65) | 19.93 |
| PE | L_V2 | Visual2 | 0.148 | < 0.001 | 0.67 (0.6-0.73) | 20.36 |
| PE | L_V3 | Visual2 | 0.121 | < 0.001 | 0.64 (0.57-0.7) | 17.39 |

Note: Brain region labels are derived from the Glasser et al., 2016 parcellation. Network affiliations are derived from Ji et al., 2019. Noise ceilings were calculated using a split-half procedure (described in the main text). The Percentage of upper Noise Ceiling refers to each brain region's representational alignment statistic divided by the upper-bound (95th percentile) noise ceiling estimate. PE; Positional Encoding

**Supplementary Table 5.** Significant representational alignment between transformer by layer and cortical regions

| **Layer** | **Region** | **Network** | **Representational alignment (r)** | $p_{FDR}$ | **Noise Ceiling Mean (5th -95th percentile)** | **Percentage of upper Noise Ceiling** |
|---|---|---|---|---|---|---|
| 1 | R_V1 | Visual1 | 0.151 | < 0.001 | 0.62 (0.54-0.69) | 21.88 |
| 1 | L_V1 | Visual1 | 0.138 | < 0.001 | 0.67 (0.59-0.73) | 18.78 |
| 1 | R_V3 | Visual2 | 0.115 | < 0.001 | 0.58 (0.5-0.65) | 17.63 |
| 2 | L_PGi | Default | 0.075 | 0.0291 | -0.0 (-0.0-0.13) | 59.00 |
| 2 | R_V1 | Visual1 | 0.276 | < 0.001 | 0.62 (0.54-0.69) | 39.99 |
| 2 | L_V1 | Visual1 | 0.271 | < 0.001 | 0.67 (0.59-0.73) | 36.88 |
| 2 | R_V2 | Visual2 | 0.266 | < 0.001 | 0.62 (0.53-0.69) | 38.42 |
| 2 | R_V3 | Visual2 | 0.22 | < 0.001 | 0.58 (0.5-0.65) | 33.73 |
| 2 | R_V4 | Visual2 | 0.086 | 0.0291 | 0.57 (0.47-0.65) | 13.21 |

| | | | | | | |
|---|---|---|---|---|---|---|
| 2 | L_V2 | Visual2 | 0.237 | < 0.001 | 0.67 (0.6-0.73) | 32.6 |
| 2 | L_V3 | Visual2 | 0.19 | < 0.001 | 0.64 (0.57-0.7) | 27.31 |
| 3 | R_6ma | Cingulo-Opercular | 0.136 | 0.0194 | 0.61 (0.52-0.69) | 19.8 |
| 3 | R_p24 | Cingulo-Opercular | 0.217 | < 0.001 | 0.21 (0.07-0.33) | 65.47 |
| 3 | L_a32pr | Cingulo-Opercular | 0.157 | 0.0111 | 0.54 (0.44-0.64) | 24.62 |
| 3 | L_p24 | Cingulo-Opercular | 0.261 | < 0.001 | 0.23 (0.1-0.34) | 76.13 |
| 3 | R_8Av | Default | 0.184 | < 0.001 | 0.47 (0.34-0.58) | 31.9 |
| 3 | R_PGi | Default | 0.149 | 0.0499 | 0.13 (0.01-0.25) | 59.15 |
| 3 | R_PGs | Default | 0.201 | 0.0111 | 0.32 (0.18-0.45) | 44.3 |
| 3 | L_8Av | Default | 0.197 | < 0.001 | 0.38 (0.25-0.5) | 39.7 |
| 3 | L_8Ad | Default | 0.154 | < 0.001 | 0.11 (-0.0-0.23) | 66.81 |
| 3 | L_PGi | Default | 0.16 | < 0.001 | -0.0 (-0.0-0.13) | 125.87 |
| 3 | L_PGs | Default | 0.234 | < 0.001 | 0.33 (0.22-0.43) | 54.84 |
| 3 | R_d32 | Frontoparietal | 0.198 | < 0.001 | 0.2 (0.08-0.31) | 63.09 |
| 3 | R_a47r | Frontoparietal | 0.149 | 0.0269 | 0.28 (0.14-0.4) | 36.85 |
| 3 | R_11l | Frontoparietal | 0.139 | 0.0269 | 0.29 (0.15-0.41) | 33.98 |
| 3 | R_s6-8 | Frontoparietal | 0.192 | < 0.001 | 0.53 (0.43-0.61) | 31.26 |
| 3 | R_TE1p | Frontoparietal | 0.17 | < 0.001 | 0.34 (0.22-0.45) | 38.13 |
| 3 | R_PFm | Frontoparietal | 0.163 | < 0.001 | 0.59 (0.49-0.67) | 24.36 |
| 3 | L_8BM | Frontoparietal | 0.12 | 0.0499 | 0.71 (0.63-0.77) | 15.68 |
| 3 | L_a47r | Frontoparietal | 0.19 | < 0.001 | 0.53 (0.43-0.62) | 30.49 |
| 3 | L_a9-46v | Frontoparietal | 0.135 | 0.0194 | 0.45 (0.32-0.56) | 24.26 |
| 3 | L_s6-8 | Frontoparietal | 0.154 | 0.0111 | 0.25 (0.09-0.39) | 39.62 |
| 3 | L_TE1p | Frontoparietal | 0.178 | < 0.001 | 0.11 (-0.0-0.24) | 74.71 |
| 3 | L_PFm | Frontoparietal | 0.188 | < 0.001 | 0.58 (0.48-0.66) | 28.31 |
| 3 | L_p47r | Frontoparietal | 0.162 | < 0.001 | 0.5 (0.39-0.6) | 26.9 |
| 3 | R_V1 | Visual1 | 0.165 | 0.0111 | 0.62 (0.54-0.69) | 23.91 |

| | | | | | | |
|---|---|---|---|---|---|---|
| 3 | L_V1 | Visual1 | 0.216 | < 0.001 | 0.67 (0.59-0.73) | 29.4 |
| 3 | R_V2 | Visual2 | 0.15 | 0.0194 | 0.62 (0.53-0.69) | 21.67 |
| 3 | L_V3 | Visual2 | 0.193 | 0.0111 | 0.64 (0.57-0.7) | 27.74 |
| 4 | R_8Av | Default | 0.21 | < 0.001 | 0.47 (0.34-0.58) | 36.41 |
| 4 | R_47l | Default | 0.26 | < 0.001 | 0.06 (-0.0-0.19) | 135.23 |
| 4 | R_PGi | Default | 0.279 | < 0.001 | 0.13 (0.01-0.25) | 110.75 |
| 4 | R_PGs | Default | 0.359 | < 0.001 | 0.32 (0.18-0.45) | 79.12 |
| 4 | L_8Av | Default | 0.177 | 0.0194 | 0.38 (0.25-0.5) | 35.67 |
| 4 | L_8BL | Default | 0.207 | 0.0332 | 0.02 (-0.0-0.15) | 137.96 |
| 4 | L_PGs | Default | 0.199 | < 0.001 | 0.33 (0.22-0.43) | 46.64 |
| 4 | R_a47r | Frontoparietal | 0.322 | < 0.001 | 0.28 (0.14-0.4) | 79.64 |
| 4 | R_11l | Frontoparietal | 0.174 | 0.0332 | 0.29 (0.15-0.41) | 42.54 |
| 4 | R_s6-8 | Frontoparietal | 0.185 | 0.0436 | 0.53 (0.43-0.61) | 30.12 |
| 4 | R_TE1p | Frontoparietal | 0.22 | < 0.001 | 0.34 (0.22-0.45) | 49.34 |
| 4 | R_p47r | Frontoparietal | 0.244 | < 0.001 | 0.13 (-0.0-0.26) | 94.57 |
| 4 | L_a47r | Frontoparietal | 0.256 | < 0.001 | 0.53 (0.43-0.62) | 41.08 |
| 4 | L_TE1p | Frontoparietal | 0.263 | < 0.001 | 0.11 (-0.0-0.24) | 110.39 |
| 4 | L_PFm | Frontoparietal | 0.173 | 0.0436 | 0.58 (0.48-0.66) | 26.05 |
| 4 | L_45 | Language | 0.189 | < 0.001 | -0.0 (-0.0-0.1) | 196.64 |

Note: Brain region labels are derived from the Glasser et al., 2016 parcellation. Network affiliations are derived from Ji et al., 2019. The Percentage of upper Noise Ceiling refers to each brain region's representational alignment statistic divided by the upper-bound (95th percentile) noise ceiling estimate.

**Supplementary Table 6: Model accuracy and correspondence to human behaviour when manipulating trained Transformer model temperature.**

| Temperature | Model Accuracy | | | Comparison to Human Accuracy | | |
|---|---|---|---|---|---|---|
| | Mean | STD | SEM | Abs difference | rho | $p^{FDR}$ |
| 0.0001 | 49.51 | 12.74 | 3.29 | -36.51 | -0.01 | 0.932 |
| 0.001 | 49.32 | 12.27 | 3.17 | -36.71 | 0.01 | 0.932 |

| | | | | | | |
|---|---|---|---|---|---|---|
| 0.01 | 51.54 | 12.36 | 3.19 | -34.31 | -0.02 | 0.932 |
| 0.1 | 71.48 | 12.75 | 3.29 | -12.77 | 0.28 | 0.005 |
| 0.25 | 90.19 | 10.6 | 2.74 | 7.43 | 0.36 | < .001 |
| 0.5 | 96.05 | 9.26 | 2.39 | 13.76 | 0.32 | 0.001 |
| 0.75 | 97.41 | 7.77 | 2.01 | 15.23 | 0.31 | 0.002 |
| 1 | 97.84 | 7.12 | 1.84 | 15.69 | 0.3 | 0.003 |
| 1.25 | 98.09 | 6.66 | 1.72 | 15.96 | 0.31 | 0.002 |
| 1.5 | 97.9 | 7.13 | 1.84 | 15.76 | 0.27 | 0.007 |
| 2 | 95.8 | 9.45 | 2.44 | 13.49 | 0.49 | < .001 |
| 3 | 83.7 | 15.87 | 4.1 | 0.43 | 0.61 | < .001 |
| 5 | 63.64 | 18.22 | 4.7 | -21.24 | 0.6 | < .001 |
| 10 | 43.7 | 8.44 | 2.18 | -42.78 | 0.47 | < .001 |
| 100 | 32.35 | 4.06 | 1.05 | -55.04 | 0.33 | 0.001 |
| 1000 | 32.16 | 3.94 | 1.02 | -55.24 | 0.32 | 0.001 |

**Supplementary Table 7.** Five top performing brain regions for each fMRI decoding analysis

| **Decoding analysis** | **Region** | **Mean Decoding (%)** | ***t*** | $p_{FDR}$ |
|---|---|---|---|---|
| Visual K-means (chance = 2.84%) | R_V1 | 4.745 | 5.865 | <.001 |
| | L_V1 | 4.63 | 7.983 | <.001 |
| | R_V2 | 4.606 | 6.033 | <.001 |
| | L_V2 | 4.375 | 5.259 | <.001 |
| | R_V4 | 4.352 | 4.565 | 0.002 |

| | | | | |
|---|---|---|---|---|
| Relational Complexity (chance = 33.33%) | R_8Av | 58.426 | 19.816 | <.001 |
| | L_PFm | 58.241 | 22.312 | <.001 |
| | R_PFm | 57.755 | 17.676 | <.001 |
| | L_8C | 56.852 | 17.681 | <.001 |
| | L_PGs | 56.667 | 16.154 | <.001 |
| Response (chance = 25%) | R_3b | 37.581 | 10.294 | <.001 |
| | R_1 | 35.507 | 8.256 | <.001 |
| | R_4 | 35.067 | 9.218 | <.001 |
| | R_3a | 32.893 | 7.507 | <.001 |
| | L_V1 | 32.836 | 7.14 | <.001 |
